# Modification of cell wall polysaccharide spatially controls cell division in *Streptococcus mutans*

**DOI:** 10.1101/2020.06.26.173716

**Authors:** Svetlana Zamakhaeva, Catherine T. Chaton, Jeffrey S. Rush, Sowmya Ajay Castro, Alexander E. Yarawsky, Andrew B. Herr, Nina M. van Sorge, Helge C. Dorfmueller, Gregory I. Frolenkov, Konstantin V. Korotkov, Natalia Korotkova

## Abstract

Bacterial cell division is driven by a tubulin homolog FtsZ, which assembles into the Z-ring structure leading to the recruitment of the cell division machinery. In ovoid-shaped Gram-positive bacteria, such as streptococci, MapZ guides Z-ring positioning at cell equators through an, as yet, unknown mechanism. The cell wall of the important dental pathogen *Streptococcus mutans* is composed of peptidoglycan decorated with Serotype *c* Carbohydrates (SCCs). Here, we show that an immature form of SCC, lacking the recently identified glycerol phosphate (GroP) modification, coordinates Z-ring positioning. Pulldown assays using *S. mutans* cell wall combined with binding affinity analysis identified the major cell separation autolysin, AtlA, as an SCC binding protein. Importantly, AtlA binding to mature SCC is attenuated due to GroP modification. Using fluorescently-labeled AtlA, we mapped SCC distribution on the streptococcal surface to reveal that GroP-deficient immature SCCs are exclusively present at the cell poles and equators. Moreover, the equatorial GroP-deficient SCCs co-localize with MapZ throughout the *S. mutans* cell cycle. Consequently, in GroP-deficient mutant bacteria, proper AtlA localization is abrogated resulting in dysregulated cellular autolysis. In addition, these mutants display morphological abnormalities associated with MapZ mislocalization leading to Z-ring misplacement. Altogether, our data support a model in which GroP-deficient immature SCCs spatially coordinate the localization of AtlA and MapZ. This mechanism ensures cell separation by AtlA at poles and Z-ring alignment with the cell equator.

**Graphical abstract:** 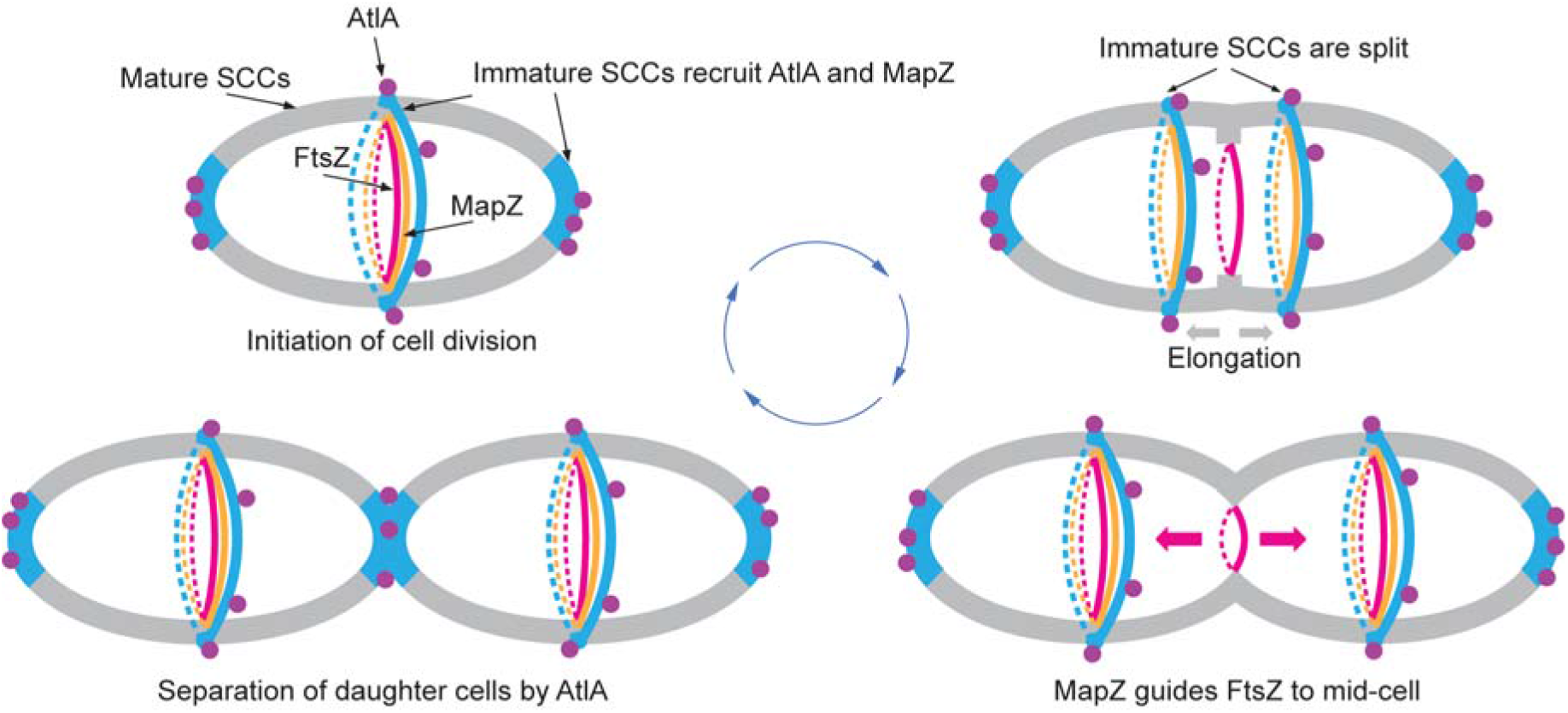

## Introduction

Bacterial cells come in a variety of shapes. The specific bacterial shapes are imposed by their cell wall, which surrounds the cytoplasmic membrane. The main structural component of the cell wall is peptidoglycan, which is composed of glycan strands that are cross-linked by penta-peptides. During cell division, new peptidoglycan is synthesized and inserted into the existing cell wall by the coordinated action of enzymes catalyzing peptidoglycan hydrolysis and synthesis. This process is tightly controlled at both the spatial and temporal level to prevent the loss of cell wall integrity and ultimately guarantee the correct cell morphology.

According to the current model, the morphogenesis of ovoid-shaped Gram-positive bacteria, such as streptococci, enterococci and lactococci, arises from a combination of septal and lateral peptidoglycan synthesis, which is coordinated by multiprotein complexes called the divisome and elongasome, respectively ^1-3^. Cell division is initiated by the recruitment and polymerization of FtsZ to form a structure called the Z-ring at mid-cell marked by a microscopically visible “wall band” or “equatorial ring” ^1,4,5^. Misplacement of FtsZ leads to severe morphological abnormalities. The Z-ring serves as a scaffold for other components of the cell division machinery, including peptidoglycan polymerases and hydrolases ^3,5^, that start cell division by synthesizing a small septal ingrowth below the equatorial ring ^1^. Early in division, new equatorial rings appear in the daughter cells, presumably due to splitting of the parental equatorial ring ^1^. During the elongation phase, the rings gradually migrate toward the equators of the daughter cells powered by two processes — splitting of the septal ingrowth and synthesis of the lateral wall ^1^. When the daughter cells have reached the size of the parental cell, and the equatorial ring approaches the mid-cell region of the nascent daughter cell, elongation is halted. At the same time, synthesis of the septal wall rapidly resumes, followed by final splitting of the complete septum by peptidoglycan hydrolases, or so-called autolysins, to allow the proper separation of the daughter cells ^1,4,5^.

It remains unclear what cues in the cell wall direct the recruitment of the cell separation autolysins. Furthermore, it is unknown how FtsZ is targeted to mid-cell. Currently, it is assumed that the correct placement of the Z-ring depends on the chromosomal origin of replication ^6^ and the FtsZ-binding protein MapZ protein MapZ ^7,8^. MapZ forms a stable protein ring that co-migrates with the equatorial ring ^7,8^, facilitating the alignment of the Z-ring perpendicular to the long axis of the cell ^6^. In Streptococcus pneumoniae, the MapZ-ring acts as a continuous guide for the orderly migration of FtsZ from the parental septum to the equatorial rings of daughter cells throughout the cell cycle ^9^. In contrast, Streptococcus mutans MapZ promotes FtsZ movement to the equators of daughter cells at a later stage in division ^10^. MapZ localization and assembly into the ring structure are reported to depend on the direct interaction of the MapZ extracellular domain with an unknown cell wall component residing in the equatorial ring ^7,11^.

The cell wall of Gram-positive bacteria contains characteristic anionic glycopolymers covalently attached to peptidoglycan. The best-studied class of cell wall glycopolymers is the canonical poly(glycerol-phosphate) and poly(ribitol-phosphate) wall teichoic acids (WTAs) expressed by *Bacillus subtilis* and *Staphylococcus aureus*, respectively ^12^. In these species, WTA-deficient mutants exhibit cell shape abnormalities, defects in the separation of daughter cells, and increased autolysis ^13-17^, indicating a functional connection between WTA and the cell division machinery. Many streptococci, including the important human dental pathogen *S. mutans*, lack canonical WTAs. Instead, they express rhamnose (Rha)-containing polysaccharides with a conserved repeating →3)α-Rha(1→2)α-Rha(1→ disaccharide backbone modified with species-specific and serotype-specific glycosyl side-chains. *S. mutans* strains are classified into four serotypes based on variations in the glycosyl side-chains, with serotype *c* being the most common in the oral cavity ^18,19^. The polysaccharide in *S. mutans* serotype *c* is referred to as serotype *c*-specific carbohydrate (SCC) and contains α-glucose (Glc) side-chains attached to the 2-position of the α-1,3 linked Rha ^20^. The homologous polysaccharide in *Streptococcus pyogenes* (Group A *Streptococcus* or GAS), referred to as the Lancefield group A carbohydrate (GAC), carries β-N-acetylglucosamine (GlcNAc) side-chain modifications attached to the 3-position of the α-1,2 linked Rha ^21,22^. There is substantial evidence that SCC and GAC are critical for cell division of streptococci ^23-27^. We recently demonstrated that SCC and GAC are also negatively charged polysaccharides, similar to WTAs, through decoration with glycerol phosphate (GroP) moieties ^28^. NMR analysis of GAC revealed that GroP is attached to the GlcNAc side-chains of GAC at the C6 hydroxyl group ^28^.

In this study, we link GroP modification of SCC to spatial regulation of streptococcal cell division. We show that structurally-diverse SCCs display a specific distribution on the *S. mutans* cell surface with cell equators and poles being populated by ‘immature SCCs’, which are deficient in the GroP modification. These immature SCCs inform the proper positioning of MapZ and the major cell separation autolysin AtlA. Thus, the presence of GroP-modified SCC in the streptococcal cell wall provides an exclusion strategy for critical cell division proteins involved in the first and the final stages of streptococcal cell division.

## Results

### GroP is attached to the Glc side-chains of SCC

We have previously shown that GroP attachment to SCC and GAC is catalyzed by a dedicated GroP transferase encoded by *sccH*/*gacH* ^28^. This gene is located in the 12-gene loci, *sccABCDEFGHMNPQ* (Fig. 1a), and *gacABCDEFGHIJKL* ^29^, which encode the biosynthesis machinery for SCC and GAC, respectively. Because in GAS, GacH is required to transfer GroP to the GlcNAc side-chains of GAC ^28^, we suggested that SccH similarly modifies the Glc side-chains of SCC with GroP in *S. mutans*. To test this hypothesis, we generated *S. mutans* c serotype strains that were devoid of the Glc side-chains. In GAS, the GtrB-type glycosyltransferase, GacI, is critical for GAC GlcNAc side-chain modification through the formation of the donor molecule GlcNAc-phosphate-undecaprenol ^30^. The SCC gene cluster (Fig. 1a) contains two putative GtrB-type glycosyltransferases encoded by *sccN* and *sccP*. To investigate the function of SccN and SccP in the synthesis of the Glc donor for side-chain addition to SCC, we deleted *sccN* and *sccP* in *S. mutans* serotype *c* strain Xc, creating Δ*sccN*, Δ*sccP*, and the double mutant Δ*sccN*Δ*sccP*. The glycosyl composition of purified SCCs exhibited a significantly reduced amount of Glc in Δ*sccN* and Δ*sccN*Δ*sccP* (Fig. 1b). In contrast, the deletion of *sccP* alone did not affect Glc levels. The Glc content of SCC was restored in Δ*sccN* by complementation with *sccN* on an expression plasmid (Δ*sccN:*p*sccN*, Fig. 1b). These data strongly support a major role for SccN in Glc side-chain formation of SCC. Interestingly, the Glc content of SCC in Δ*sccN*Δ*sccP* was lower than in Δ*sccN* (Fig. 1b), suggesting that *sccP* might play a minor role in providing a Glc donor for modification of SCC with the side-chains (Supplementary Fig. 1a). Furthermore, we found that the glycerol and phosphate content in the polysaccharide isolated from Δ*sccN* was significantly reduced (Fig. 1b), similar to the Δ*sccH* mutant (Fig. 1b). We have previously provided conclusive evidence that glycerol and phosphate detected in this analysis are, in fact, GroP ^28^. The deficiency of glycerol and phosphate in the Δ*sccN* mutant was reversed by complementation with *sccN*, supporting the conclusion that the Glc-side chains of SCC are further modified with GroP by SccH (Supplementary Fig. 1a, b and c).

**Fig. 1.**
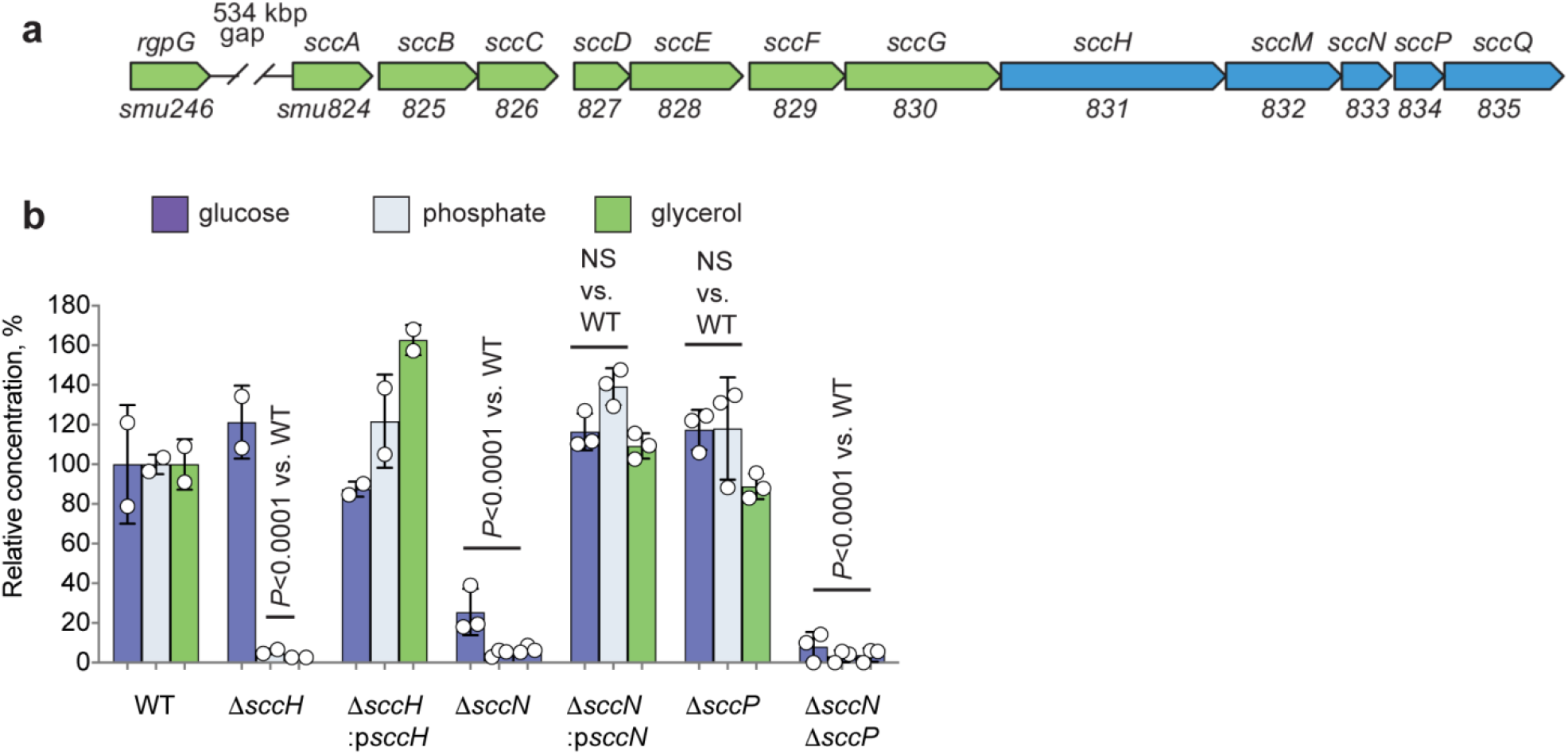
Modification of the SCC polyrhamnose backbone with Glc side-chains and GroP. (**a**) Schematic representation of the SCC biosynthetic gene cluster. SCC gene cluster *smu.*824-835 was renamed *sccABCDEFGHMNPQ* ^28^. (**b**) Composition analysis of SCCs isolated from *S. mutans* WT, Δ*sccH*, Δ*sccH:psccH*, Δ*sccN*, Δ*sccN:psccN*, Δ*sccP*, and Δ*sccN*Δ*sccP*. The concentrations ofGlc, phosphate, and glycerol are normalized to Rha content and presented as a percentage of the ratios in the WT strain. Glc was analyzed by GC-MS. Phosphate and glycerol were analyzed by colorimetric assays. Columns and error bars represent the mean and S.D., respectively (biologically independent replicates for Δ*sccN*, Δ*sccN:psccN*, Δ*sccP*, and Δ*sccN*Δ*sccP* were three, and for WT, Δ*sccH*, Δ*sccH:psccH* were two). *P* values were calculated by 2-way ANOVA with Bonferroni’s multiple comparison test.

### GroP modification controls the self-aggregation and morphology of *S. mutans*

We observed that planktonic Δ*sccH* and Δ*sccN* have a strong tendency to spontaneously sediment after overnight growth, as compared to the wild type (WT) strain and the complemented strains, Δ*sccH*:p*sccH* and Δ*sccN*:p*sccN*, which remained in suspension (Fig. 2a). Microscopic analysis of bacteria revealed that Δ*sccH* and Δ*sccN*, but not the WT strain, Δ*sccH*:p*sccH* and Δ*sccN*:p*sccN*, formed the typical short chains that clump together (Fig. 2b). Furthermore, the bacterial aggregates of Δ*sccH* and Δ*sccN* were not dispersed when DNAse was added to the growth medium (Supplementary Fig. 2), suggesting that this bacterial behavior is due to cell-cell interactions.

**Fig. 2.**
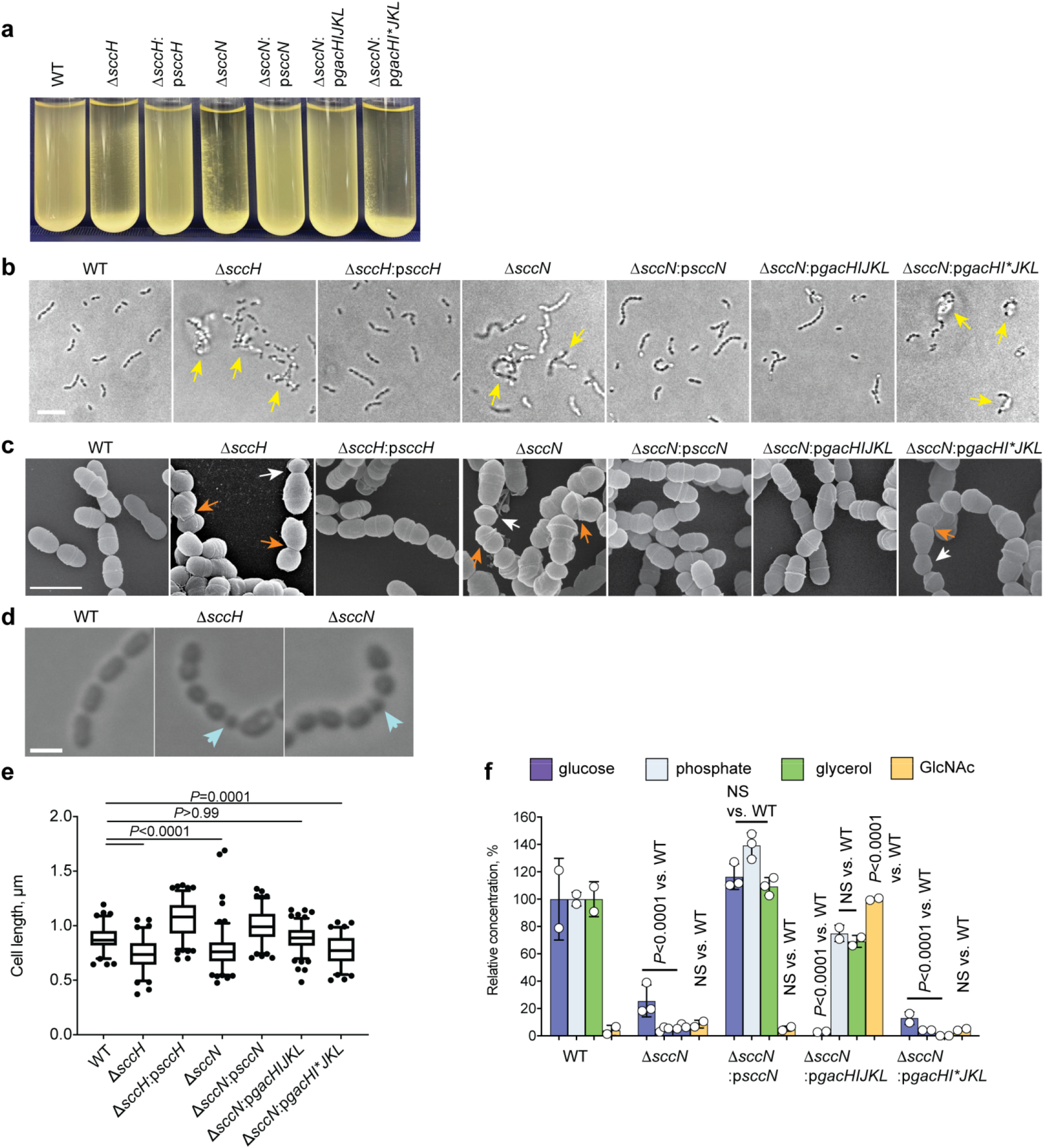
GroP modification of SCC controls the cell aggregation and morphology of *S. mutans*. (**a**) Sedimentation phenotype of *S. mutans* WT and specific mutants after overnight growth in THY broth. (**b**) Differential interference contrast (DIC) images of bacteria grown in THY broth overnight. Yellow arrows indicate bacterial aggregates. Scale bar is 5 µm. (**c** and **d**) Morphological defects of *S. mutans* deficient in GroP and glycosyl side-chain decorations. (**c**) Scanning electron micrographs of *S. mutans* WT and specific mutants. Exponentially growing bacteria were fixed, dehydrated stepwise, and viewed by scanning electron microscopy. White and orange arrows denote small round cells and the cells with skewed division planes, respectively. (**d**) DIC images of exponentially growing bacteria. Blue arrows denote small round cells. Representative images from at least three independent experiments are shown in **a, b, c** and **d**. Scale bar is 1 µm in **c** and **d**. (**e**) Cell length of bacterial strains at the mid-exponential phase. DIC images of bacterial cells were analyzed using the software ImageJ to quantify cell length and width. *P*-values were determined by one-way ANOVA for on ranks with Dunn’s test. (**f**) Composition analysis of polysaccharides isolated from *S. mutans* WT, Δ*sccN*, Δ*sccN:psccN*, Δ*sccN:pgacHIJKL* and Δ*sccN:pgacHI*JKL*. The concentrations of Glc, phosphate, and glycerol are normalized to Rha content and presented relative to the WT strain. The concentration of GlcNAc is normalized to Rha content and presented relative to Δ*sccN*:p*gacHIJKL*. Glc was analyzed by GC-MS. Phosphate, glycerol, and GlcNAc were analyzed by colorimetric assays. Results of GlcNAc analysis were confirmed by GC-MS. Columns and error bars represent the mean and S.D., respectively (n = 2 biologically independent replicates were for Δ*sccN:pgacHIJKL* and Δ*sccN:pgacHI*JKL*, and GlcNAc content in all strains). Glc, phosphate, and glycerol contents in *S. mutans* WT, Δ*sccN*, and Δ*sccN:psccN* as in Fig 1 b. *P* values were calculated by 2-way ANOVA with Bonferroni’s multiple comparison test.

Analysis of WT, Δ*sccH*, and Δ*sccN* by scanning electron microscopy (SEM) and differential interference contrast (DIC) microscopy revealed that the mutant cells have severe cell division defects. The WT bacteria displayed the characteristic oval shape with the average cell length of 0.88±0.11 µm in the exponential growth phase (Fig. 2c, d, and e and Supplementary Table 3). In contrast, the majority of the Δ*sccH* and Δ*sccN* cells were significantly shorter than the WT cells with the average length of 0.74±0.14 µm and 0.78±0.18 µm, respectively (Fig. 2e and Supplementary Table 3). Additionally, the distribution in the length of the individual Δ*sccH* and Δ*sccN* cells differed significantly compared to WT (Supplementary Fig. 3); 21% of the Δ*sccH* cells (n = 86) and 14% of the Δ*sccN* cells (n = 117) were abnormally small, while only 2% of the WT cells displayed the minimal cell length (Supplementary Table 3). The small cells of Δ*sccH* and Δ*sccN* were frequently paired with larger cells (Fig. 2c and d), implying that these cells resulted from asymmetrically cell division, giving rise to daughter cells with unequal sizes. Finally, cells with the orientation of the division plane not perpendicular to the long axis of the cell were also observed in the Δ*sccH* and Δ*sccN* mutants (Fig. 2c). The morphological phenotypes were restored to WT in Δ*sccH*:p*sccH* and Δ*sccN*:p*sccN* (Fig. 2c, e and Supplementary Table 3). These phenotypes were also correlated with a significant decrease in cell viability of Δ*sccH* determined by colony-forming unit count, in comparison to WT and Δ*sccH*:p*sccH* (Supplementary Fig. 4). We thus concluded that either GroP or the epitope presented by the Glc side-chains modified with GroP are required for proper morphogenesis of *S. mutans*.

To dissect the structural requirements underlying these morphological processes, we replaced the SCC Glc side-chains with GlcNAc in *S. mutans* by plasmid-based expression of *gacHIJKL* ^30^ in Δ*sccN* creating Δ*sccN:pgacHIJKL* (Supplementary Fig.1d and e). These genes (Supplementary Fig. 1e) are required for the formation and addition of the GAC GlcNAc side-chains and GroP in GAS ^30^. Additionally, as a negative control, we expressed *gacHI*JKL* in the Δ*sccN* genetic background creating Δ*sccN:pgacHI*JKL*. This strain does not synthesize the GlcNAc donor for side-chain addition since it carries a stop codon in *gacI.* The cell wall polysaccharide purified from Δ*sccN:pgacHIJKL* showed increased levels of GlcNAc and GroP (Fig. 2f), whereas GlcNAc and GroP were not restored by expression of *gacHI*JKL* in Δ*sccN* (Fig. 2f). The morphological phenotypes of Δ*sccN* were only reversed to WT by expression of *gacHIJKL* but not *gacHI*JKL* (Fig. 2a, b, c, e, Supplementary Fig.3 and Supplementary Table 3). These observations indicate that the underlying molecular mechanisms for clumping behavior and the morphological abnormalities are GroP-dependent and independent of the specific glycosyl side chain. The simple explanation for the self-aggregation of bacteria is that Δ*sccH* and Δ*sccN* lack a negative surface charge provided by GroP, which normally contributes to electrostatic repulsion between the WT cells.

### GroP modification protects *S. mutans* from autolysis

It has been reported that the WTA-deficient mutant of *S. aureus* show increased fragility and autolysis due to mislocalization and increased abundance of the major cell division autolysin, Atl, on the bacterial surface ^15^. Hence, to understand whether and how GroP and the Glc side-chains in SCC affect *S. mutans* autolysis, we compared autolysis of the WT, Δ*sccH*, Δ*sccN* and the complemented strains Δ*sccH*:p*sccH*, Δ*sccN*:p*sccN*, Δ*sccN:pgacHIJKL* and Δ*sccN:pgacHI*JKL* by analyzing the change in OD_600_ in liquid cultures after the addition of a mild detergent Triton-X100. While *S. mutans* WT is relatively resistant to detergent-induced lysis (Fig. 3a), Δ*sccH* and Δ*sccN* are sensitive to the autolytic effect of Triton X-100. Cellular lysis was more pronounced in Δ*sccN*, than in Δ*sccH*, resulting in significant loss of turbidity of the mutant suspension after 2 hours (Fig. 3a). The phenotypes of Δ*sccH* and Δ*sccN* were reversed to WT in Δ*sccH*:p*sccH* and Δ*sccN*:p*sccN.* Furthermore, expression of *gacHIJKL*, but not *gacHI*JKL*, in the Δ*sccN* mutant restored the resistance of bacteria to autolysis. These data indicate the importance of GroP and side-chain modifications of SCC in protecting *S. mutans* from adventitious cellular lysis by an unknown autolytic enzyme.

**Fig. 3.**
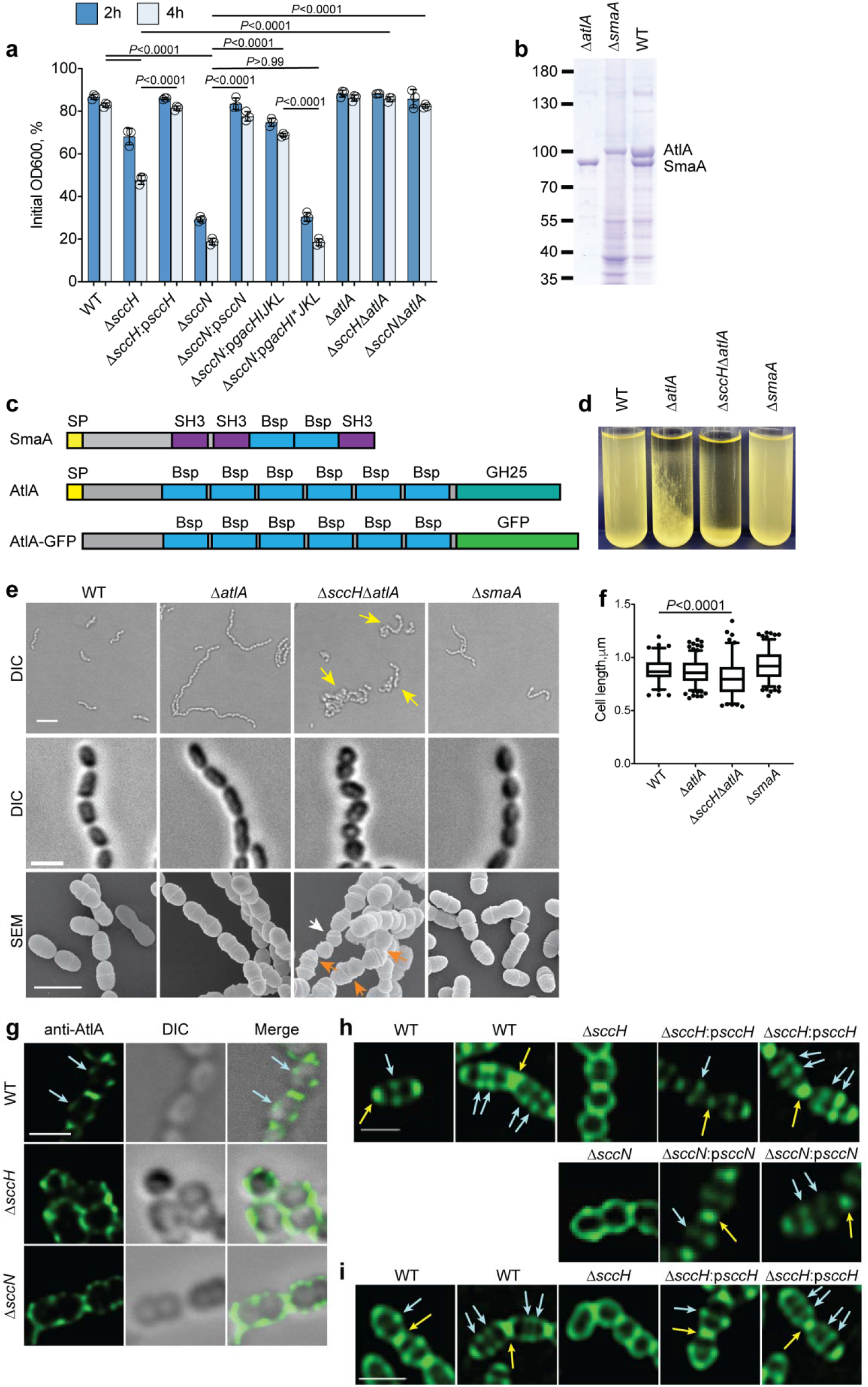
AtlA mediates autolysis of the *S. mutans* mutants lacking GroP and the side-chain in SCC. (**a**) The autolytic activity of *S. mutans* WT, Δ*sccH*, Δ*sccH:psccH*, Δ*sccN*, Δ*sccN:psccN*, Δ*sccN:pgacHIJKL* Δ*sccN:pgacHI*JKL*, Δ*atlA*, Δ*sccH*Δ*atlA* and Δ*sccN*Δ*atlA.* Exponentially growing strains were allowed to autolyze in 0.1% Triton X-100. The autolysis was monitored after 2 and 4 h as the decrease in OD_600_. Results were normalized to the OD_600_ at time zero (OD_600_ of 0.5). Columns and error bars represent the mean and S.D., respectively (n = 3 biologically independent replicates). *P*-values were determined by two-way ANOVA with Tukey’s multiple comparisons test. (**b**) A pulldown of WT, Δ*atlA*, and Δ*smaA* cell surface proteins using the cell wall purified from *S. mutans*. AtlA and SmaA were identified as proteins bound to the cell wall. Cell surface proteins were extracted from WT, Δ*atlA*, and Δ*smaA* with 8 M urea, refolded overnight, and incubated with the lyophilized cell wall purified from *S. mutans* WT. Bound proteins were extracted with SDS sample buffer and separated on 10% SDS-PAGE. Representative image from at least three independent experiments is shown. (**c**) Diagram of the domain structure of SmaA, AtlA and AtlA-GFP. SP (yellow box) denotes signal peptide. (**d**) The growth phenotype of WT, Δ*atlA*, Δ*sccH*Δ*atlA* and Δ*smaA.* Bacteria were grown in THY broth overnight. (**e**) Differential interference contrast (DIC) images of bacteria grown in THY broth overnight (top panels, scale bar is 5 µm) and exponentially growing bacteria (middle panels, scale bar is 1 µm). Yellow arrows indicate bacterial aggregates. Scanning electron micrographs of WT, Δ*atlA*, Δ*sccH*Δ*atlA* and Δ*smaA* (bottom panels, scale bar is 1 µm). Exponentially growing bacteria were fixed, dehydrated stepwise, and viewed by scanning electron microscopy. White and orange arrows denote small round cells and the cells with skewed division plane, respectively. (**f**) Cell length of bacterial strains at the mid-exponential phase. DIC images of bacterial cells were analyzed using the software ImageJ to quantify cell length and width. *P*-values were determined by one-way ANOVA for on ranks with Dunn’s test. (**g**) Immunolocalization of AtlA on the surface of WT, Δ*sccH*, and Δ*sccN* cells. Cells were grown to mid-log phase, immunostained with anti-AtlA antibodies, and examined by differential interference contrast (DIC) (center panels) and fluorescence microscopy (left panels). An overlay of AtlA immunostaining and DIC is shown in the right panels. Blue arrows depict equatorial sites labeled by AtlA-GFP. (**h**) Binding of AtlA-GFP to exponentially growing WT, Δ*sccH*, Δ*sccH*:p*sccH*, Δ*sccN* and Δ*sccN*:p*sccN* cells. Yellow and blue arrows show polar and equatorial sites, respectively, labeled by AtlA-GFP. (**i**) Binding of AtlA-GFP to sacculi of WT, Δ*sccH* and Δ*sccH*:p*sccH*. Scale bar is 1 µm in **g, h**, and **i**. The experiments depicted in **b, d, e, g, h**, and **i** were performed independently three times and yielded the same results.

### AtlA promotes autolysis of the mutants lacking GroP and side-chain modifications

A possible mechanistic explanation for enhanced autolysis of Δ*sccH* and Δ*sccN* could be that the absence of GroP and Glc side-chain modifications in SCC causes a dysregulated localization of an unknown autolysin. To identify autolytic enzymes that interact with SCCs, we stripped the cell surface-associated proteins from *S. mutans* WT cells using 8 M urea, re-folded the released proteins by dialysis, and used the re-folded proteins for pulldown experiments with the protein-deficient cell wall material purified from *S. mutans*. Two major proteins, isolated by the pulldown approach, were identified as AtlA and SmaA by LC-MS/MS analysis (Fig. 3b). One of these, AtlA, is the major autolysin involved in the separation of daughter cells after division ^31-34^. The protein contains an N-terminal putative cell wall-binding domain, which consists of six Bsp repeats (PF08481) and a C-terminal family GH25 catalytic domain (PF01183) (Fig. 3c). SmaA has two Bsp repeats and three bacterial Src homology 3 (SH3) domains (Fig. 3c). Although SmaA has been implicated in peptidoglycan hydrolysis ^35^, the protein lacks a putative catalytic domain.

To confirm the identities of the proteins, we generated Δ*atlA* and Δ*smaA* deletion mutants. Using a similar approach as described above, surface proteins were stripped from the Δ*atlA* and Δ*smaA* bacteria and used in pulldown experiments. In agreement with the results of protein identification, the bands corresponding to AtlA and SmaA were absent in Δ*atlA* and Δ*smaA*, respectively (Fig. 3b). To investigate the contribution of AtlA and SmaA in Triton X-100-induced autolysis of WT, Δ*sccH*, and Δ*sccN*, we deleted *atlA* in the Δ*sccH* and Δ*sccN* backgrounds, and *smaA* in the Δ*atlA* and Δ*sccN* backgrounds, generating the Δ*smaA*Δ*atlA*, Δ*sccH*Δ*atlA*, Δ*sccN*Δ*atlA*, and Δ*sccN*Δ*smaA* mutants. As expected, the Δ*atlA*, Δ*smaA*, and Δ*smaA*Δ*atlA* mutants are relatively resistant to detergent-stimulated lysis (Fig. 3a, Supplementary Fig. 5). The Δ*sccN*Δ*smaA* mutant displayed increased cellular lysis, similar to Δ*sccN*, indicating that SmaA is not involved in detergent-induced autolysis of Δ*sccN* (Supplementary Fig. 5b). In contrast, the deletion of *atlA* in the Δ*sccH* and Δ*sccN* backgrounds restored the bacterial resistance to the autolytic effect of Triton X-100 (Fig. 3a). These results clearly show that autolysis of Δ*sccH* and Δ*sccN* is AtlA-mediated.

Next, we investigated whether the deletion of SmaA or AtlA affected morphology and self-aggregation of *S. mutans*. The Δ*smaA* cells remained in suspension after overnight growth (Fig. 3d), and showed WT cell morphology, cell separation, and cell length distribution (Fig. 3e, f, Supplementary Fig. 3 and Supplementary Table 3). The Δ*atlA* cells grew in long chains (Fig. 3e) that settle at the bottom of the tube (Fig. 3d), which is in line with the published observation ^33^. However, in contrast to Δ*sccH* and Δ*sccN*, the individual Δ*atlA* cells displayed normal dimensions, and the long chains of Δ*atlA* did not self-aggregate (Fig. 3e, f, Supplementary Fig. 3, and Supplementary Table 3). Finally, we examined the phenotype of the Δ*sccH*Δ*atlA* mutant to understand more about the link between AtlA and GroP modification of SCC. The phenotype of Δ*sccH*Δ*atlA* was a combination of the phenotypes of single mutants. Notably, similar to Δ*atlA*, the double mutant displayed a chaining phenotype, and like Δ*sccH*, the long chains of the double mutant formed clumps (Fig. 3e), supporting the hypothesis that self-aggregation of Δ*sccH*Δ*atlA* is due to loss of electrostatic repulsion between the cells. Furthermore, SEM and DIC analyses of Δ*sccH*Δ*atlA* revealed a variety of aberrant cell shapes and size that contrast with the morphology of WT and Δ*atlA*, but similar to Δ*sccH*. (Fig. 3e, f, Supplementary Fig. 3 and Supplementary Table 3). Altogether, these data indicate that, although autolysis of Δ*sccH* and Δ*sccN* requires AtlA, the cell shape abnormalities observed in Δ*sccH* and Δ*sccN* are not linked to either SmaA or AtlA.

### GroP modification coordinates the localization of AtlA at the cell pole and mid-cell

Considering the established function of AtlA in the autolysis of Δ*sccH* and Δ*sccN*, we hypothesized that the enzyme is either mislocalized or over-expressed in these mutants. To examine the expression of AtlA, the protein was extracted from the cell surface of the WT, Δ*sccH*, and Δ*sccN* bacteria. Western blot analysis with anti-AtlA antibodies revealed no significant differences in the amount of AtlA recovered from *S. mutans* WT or the mutants (Supplementary Fig. 6).

Next, we monitored the localization of AtlA on the cell surface of WT, Δ*sccH*, and Δ*sccN* by immunofluorescent microscopy using anti-AtlA antibodies and fluorescent secondary antibodies (Fig. 3g). The autolysin localized to the cell poles and the mid-cell of the WT strain. Secondary antibodies alone showed no specific binding to *S. mutans* (Supplementary Fig. 7a). In the case of Δ*sccH* and Δ*sccN*, the autolysin was evenly distributed over the cell surface, and showed no distinct surface localization pattern (Fig. 3g).

To provide a more detailed picture of the bacterial regions targeted by AtlA, we fused the N-terminal Bsp repeat domain of AtlA with a green fluorescent protein (GFP) ^36^, generating AtlA-GFP (Fig. 3c). The fusion protein was added exogenously to exponentially growing *S. mutans* WT, Δ*sccH*, Δ*sccN*, Δ*sccH*:p*sccH* and Δ*sccN*:p*sccN* cells. Fluorescence microscopy imaging indicated that in newborn WT cells, AtlA-GFP predominantly targeted poles and mid-cell zones (Fig. 3h). Surprisingly, in the cells that were beginning to elongate, the sites labeled by incubation with AtlA-GFP split as a pair of rings (Fig. 3h). Splitting of the mid-cell sites was observed in 47% of the cells (64/135). The mid-cell localization of AtlA-GFP in newborn cells and the duplicated AtlA-GFP localization signal in elongating cells correlate with the position of the equatorial ring in ovococci ^4,37^. In contrast, AtlA-GFP added to Δ*sccH* and Δ*sccN* was uniformly distributed along the bacterial cell surface (Fig. 3h). The complemented strains Δ*sccH*:p*sccH* and Δ*sccN*:p*sccN* demonstrated localization of AtlA-GFP similar to the WT strain (Fig. 3h). GFP alone failed to bind to the bacteria (Supplementary Fig. 7b). Together our findings provide three important insights: i) the N-terminal Bsp repeat domain of AtlA is sufficient for proper binding of AtlA to the bacterial surface; ii) the Bsp repeat domain recognizes the specific cell wall component incorporated in cell poles and the regions corresponding to equatorial rings; iii) the absence of GroP and side-chain decorations on SCC correlate with mislocalized binding of AtlA across the entire surface of *S. mutans*.

### AtlA binds to SCC

To identify the cell surface structure targeted by AtlA-GFP, we examined the binding of the protein fusion to sacculi purified from *S. mutans* WT, Δ*sccH* and Δ*sccH*:p*sccH*. The sacculi were free of LTA, proteins, lipids and nucleic acids ^30^. Fluorescence microscopy imaging revealed that AtlA-GFP associated with the sacculi derived from these cells with patterns very similar to that observed with intact *S. mutans* cells (Fig. 3i). AtlA-GFP was found primarily at the poles and mid-cell sites of WT and Δ*sccH*:p*sccH* sacculi, but was distributed evenly along the surface of Δ*sccH* sacculi. Furthermore, the splitting of AtlA-GFP-labeled mid-cell sites in WT and the complemented cells was observed. Splitting was detected in 50% of WT cells (63/125). GFP alone was unable to attach to the sacculi (Supplementary Fig. 7c). This observation confirmed that the N-terminal domain of AtlA recognizes a cell wall component, either SCC or peptidoglycan, associated with the cell equators and poles.

Notably, secreted proteins consisting of multiple Bsp repeats are widespread in Firmicutes, with most bacteria belonging to the Streptococcus genus (Supplementary Fig. 8). Intriguingly, there is an obvious correlation between the co-occurrence of the genes encoding proteins with Bsp repeats and Rha-containing cell wall polysaccharides in these bacteria (Supplementary Table 4). In fact, in many streptococci, a gene encoding a putative autolysin with Bsp repeats is situated immediately downstream of the gene loci involved in the biosynthesis of the Rha-containing polysaccharide, raising the possibility that the Bsp repeat domain binds Rha-containing cell wall polysaccharides. Thus, to examine the role of SCCs in the recruitment of AtlA, we applied a co-sedimentation assay that exploits the property of AtlA-GFP to associate with the cell wall material purified from WT *S. mutans*, Δ*sccH* and Δ*sccH*:p*sccH*. In agreement with the fluorescent microscopy experiments, we observed a very strong binding of the fusion protein to WT *S. mutans*, Δ*sccH* and Δ*sccH*:p*sccH* cell walls (Fig. 4a). GFP alone did not attach to cell walls purified from *S. mutans* WT (Fig. 4b). Next, we chemically cleaved the polysaccharides from peptidoglycan using mild acid hydrolysis. SCCs were efficiently released from the cell walls in these conditions (Supplementary Fig. 9a). We observed a significant reduction in AtlA-GFP binding to the SCC-depleted cell wall (Fig. 4a), supporting the role of SCCs in targeting of AtlA to the equatorial rings and the poles of the WT bacteria. To further investigate the participation of SCCs in cellular localization of AtlA, we constructed a SCC-deficient mutant by in-frame deletion of the *rgpG* gene whose product catalyzes the first step in SCC biosynthesis ^38^. The Δ*rgpG* mutant was devoid of polyrhamnose polysaccharide (<0.5% WT level), and in agreement with the published data ^39,40^, demonstrated self-aggregation and aberrant morphology (Supplementary Fig. 9a and b). Applying AtlA-GFP to the intact Δ*rgpG* bacteria (Supplementary Fig. 9c) and to the cell wall purified from Δ*rgpG* (Fig. 4b), no evidence of binding was observed in either assay. These results strongly indicate that SCC is a binding receptor of AtlA.

**Fig. 4.**
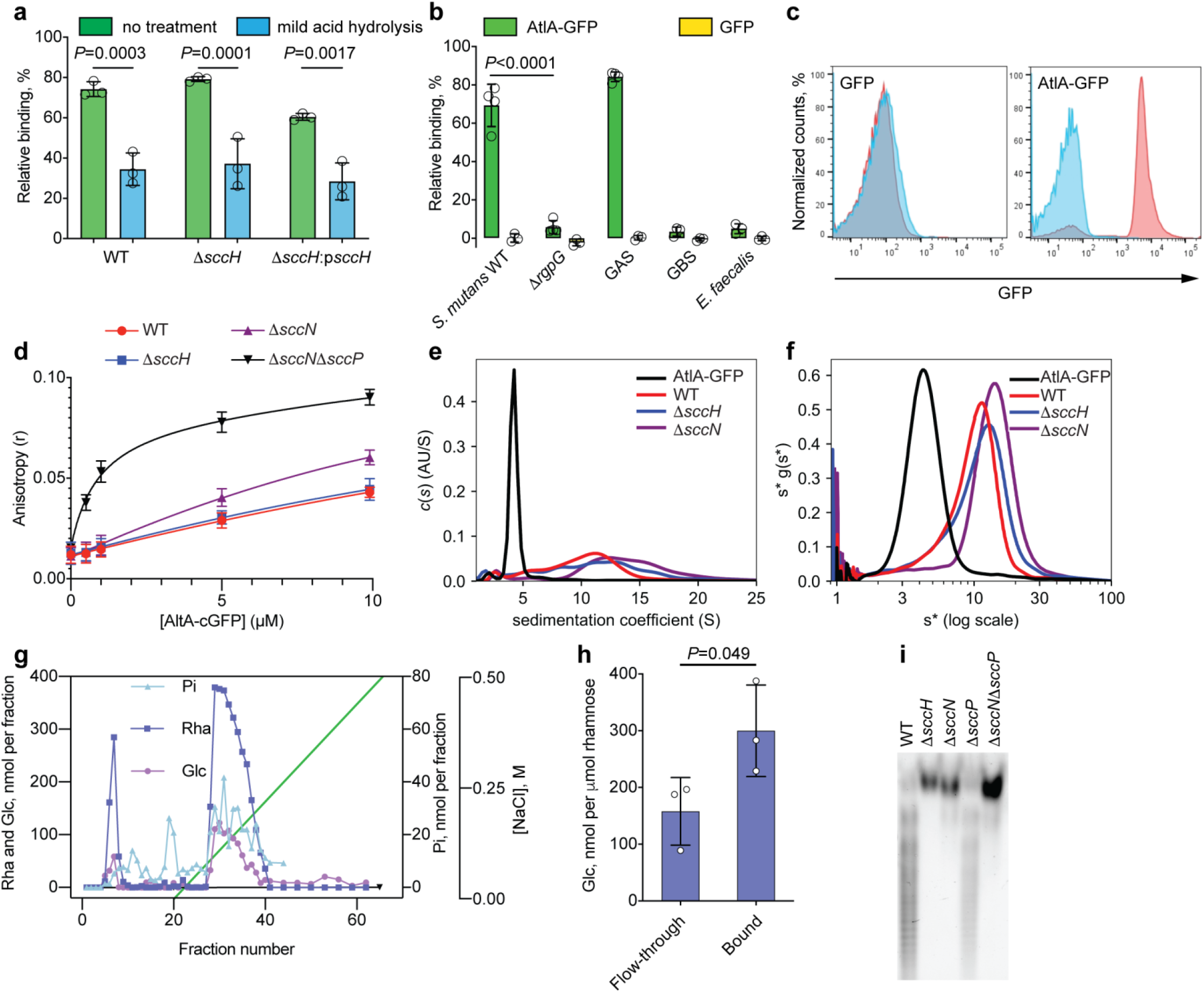
AtlA-GFP binds to the polyrhamnose backbone of SCC. (**a**) Binding of AtlA-GFP to *S. mutans* cell wall requires intact SCC. Co-sedimentation assay of AtlA-GFP with cell walls purified from WT, Δ*sccH* and Δ*sccH*:p*sccH.* The protein fusion was incubated with intact cell walls (no treatment) or the cell walls subjected to mild acid hydrolysis to cleave SCCs from peptidoglycan (mild acid hydrolysis). The efficiency of SCC release from cell walls is shown in Supplementary Figure 9a. (**b**) Co-sedimentation assay of AtlA-GFP and GFP with cell walls purified from *S. mutans* WT, Δ*rgpG*, GAS, *S. equi*, GBS and *E. faecalis*. Data in **a** and **b** are presented as a percentage of fluorescence of the pellet normalized to the total fluorescence of the sample. Columns and error bars represent the mean and S.D., respectively (n = 3 biologically independent replicates). *P* values were calculated by a two-way ANOVA with Tukey’s multiple comparisons test. (**c**) AtlA-GFP binds to *E. coli* expressing polyrhamnose on the cell surface. Flow cytometry analysis of AtlA-GFP (right panel) and GFP (left panel) binding to the polyrhamnose-expressing *E. coli* (red) and its parental strain (blue). For flow cytometric analysis, at least 10000 events were collected. Experiments depicted in **c** were performed independently three times and yielded the same results. Histograms of representative results are shown. (**d**) Fluorescence anisotropy of 0.5 µM ANDS-labeled SCC variants incubated with titrations of AtlA-cGFP. Binding of the Δ*sccN*Δ*sccP* SCC to AtlA-GFP was fitted to a one-site total-binding model. Error bars represent the 95% confidence interval of three independent replicates. (**e** and **f**) Analytical ultracentrifugation analysis of AtlA-GFP binding to SCCs extracted from WT, Δ*sccH* and Δ*sccN*. (**e**) The continuous c(s) distributions for 2.5 µM AtlA-GFP combined with 25 µM SCC variants. (**f**) Plot of s*g(s*) vs. log(s*) using wide distribution analysis in SedAnal. Experiments depicted in **e** and **f** were performed independently twice and yielded the same results. (**g**) Ion exchange chromatography of SCCs purified from WT *S. mutans*. SCC material was loaded onto Toyopearl DEAE-650M and eluted with a NaCl gradient (0-0.5 M). Fractions were analyzed for Rha and Glc contents by anthrone assay and total phosphate (Pi) content by malachite green assay following digestion with perchloric acid. The experiments were performed at least three times and yielded the same results. Data from one representative experiment are shown. (**h**) Composition analysis of minor and major fractions from **g**. Fractions were pooled, concentrated, and desalted by spin column and analyzed by GC-MS to determine the Rha and Glc concentrations. The concentration of Glc is presented relative to the Rha concentration. Columns and error bars represent the mean and S.D., respectively (n = 3 biologically independent replicates). *P* values were calculated by a two-tailed t-test. (**i**) SDS-PAGE analysis of ANDS-labeled SCCs extracted from *S. mutans* mutants. Representative image from at least three independent experiments is shown.

### AtlA binds to SCC via the polyrhamnose backbone

The ability of AtlA to target distinct sites on the *S. mutans* cell might be explained by binding of the autolysin to a specific form of SCC present in the equatorial rings and the poles. Our observation that AtlA-GFP is uniformly distributed over the cell surface of the side-chain-deficient mutant, Δ*sccN*, suggests that the N-terminal domain of AtlA binds to polyrhamnose regions on SCC. To determine if AtlA specifically binds the polyrhamnose backbone of SCC we compared the binding of AtlA-GFP to intact cells, or purified cell walls, of bacterial strains that express cell wall polysaccharides containing this backbone feature (GAS and *Streptococcus equi*) ^29^ with other bacterial strains that express Rha-containing cell wall polysaccharides lacking this structure (Group B Streptococcus or GBS, and *Enterococcus faecalis*) (Supplementary Fig. 10a, b, c, d and e). No significant binding was observed to the cell walls or intact cells of GBS and *E. faecalis* (Fig. 4b and Supplementary Fig. 10f). In contrast, as compared to GFP control, AtlA-GFP strongly binds to the cell wall material purified from GAS (Fig. 4b) and the intact GAS and *S. equi* cells (Supplementary Fig. 10f), suggesting that the polyrhamnose backbone of GAC, SCC and the *S. equi* cell wall polysaccharide is recognized by AtlA.

To gather additional evidence that AtlA interacts with the polyrhamnose backbone, we took advantage of a previously developed heterologous expression model in *E. coli*. Specifically, complementation of the SCC polyrhamnose biosynthetic genes *sccABCDEFG* in *E. coli* results in the decoration of lipopolysaccharide with polyrhamnose ^41,42^. The expression of polyrhamnose on the surface of *E. coli* was confirmed by anti-GAC antibodies that recognize the GAC polyrhamnose backbone (Supplementary Fig. 11). Flow cytometric analysis revealed that AtlA-GFP only bound to polyrhamnose-producing *E. coli* and not to the parental strain (Fig. 4c). These data unambiguously demonstrate that the SCC polyrhamnose backbone is sufficient to confer AtlA binding. GFP alone did not bind to the recombinant bacteria (Fig. 4c).

### GroP and the Glc side-chains hinder AtlA-GFP binding to the polyrhamnose backbone

To investigate how the presence of the SCC side-chain substituents affects recognition of the polyrhamnose backbone by AtlA, we compared AtlA binding affinities to various SCCs using fluorescence polarization anisotropy. The SCC variants prepared from WT, Δ*sccH*, Δ*sccN*, and Δ*sccN*Δ*sccP* were conjugated with a fluorescent tag (ANDS). The colorless AtlA-GFP variant protein, AtlA-cGFP, was incubated with the fluorescently labeled SCCs at varying concentrations (Fig. 4d). We observed that the binding of the Δ*sccN*Δ*sccP* SCC is the strongest, with an apparent K_d_ = 0.9 µM (95% confidence interval: 0.6-1.2), which is in the range of the binding affinity of known lectins for complex glycans ^43,44^. While binding of the WT, Δ*sccH*, and Δ*sccN* SCCs was unable to reach saturation due to solubility constraints, they all demonstrate clear evidence of binding to AtlA-cGFP (Fig. 4d). The Δ*sccN* SCC has the second-highest affinity, estimated to be at least 12-fold weaker than Δ*sccN*Δ*sccP*, with the Δ*sccH* and WT SCCs being indistinguishable under accessible conditions. Both SCCs have at least a 25-fold lower affinity compared to SCC isolated from Δ*sccN*Δ*sccP* bacteria. The differences in binding of the SCC variants suggest that the primary recognition site of the AtlA Bsp repeat domain is unmodified polyrhamnose, and the addition of branching structures decreases binding affinity, presumably due to steric hindrance. The differences in binding between the Δ*sccN*Δ*sccP* and Δ*sccN* SCCs further reinforces our hypothesis that both SccN and SccP provide the Glc donor for modification of SCC with the side-chains.

To further explore differences in binding of AtlA-GFP to the SCCs extracted from WT, Δ*sccH*, and Δ*sccN*, we employed analytical ultracentrifugation. Continuous c(s) distribution analysis estimated AtlA-GFP to have a molecular weight of ∼110 kDa, confirming that the protein fusion remains mostly monomeric at the concentration range being studied (Fig. 4e). This verifies that assembly is polysaccharide-dependent and not a function of the GFP tag. Upon addition of the WT, Δ*sccH*, or Δ*sccN* SCCs, higher-order complexation was observed between 10-20 S (Fig. 4e). The width of the distribution and the high s values of the largest species imply that many different permutations are occurring simultaneously. There is likely an equilibrium of different combinations: one AtlA protein bound to multiple SCCs, multiple proteins bound to a single SCC, and higher-order configurations of SCCs and AtlA proteins bridging and/or daisy-chaining together to form very large assemblies. Because c(s) is a weight-averaged model, molecular weight estimates are not meaningful for these types of distributions. However, the experiments demonstrate that WT SCC produces, on average, a lower-sized complex than either the Δ*sccH* or Δ*sccN* polysaccharides.

To better understand overall complexation, the same data were analyzed using wide distribution analysis (Fig. 4f). Each sample resolved into a more Gaussian distribution, with a weight-averaged sedimentation coefficient of 5.5 S for AtlA-GFP alone, 10.6 S for the WT SCC, 12.3 S for the Δ*sccH* SCC, and 14.5 S for the Δ*sccN* SCC. Based on these values, wide distribution analysis also confirms the formation of smaller complexes in the presence of WT SCCs compared to the Δ*sccH* and Δ*sccN* polysaccharides. Thus, our results indicate that AtlA-GFP binds to the polyrhamnose backbone of SCC, and the modification of SCC with GroP and the Glc side-chains obstruct AtlA-GFP attachment to the polyrhamnose backbone.

### SCC is highly heterogeneous with regard to GroP modification

AtlA localization studies together with the analysis of AtlA-GFP binding affinity are consistent with the idea that *S. mutans* equatorial sites and poles contain SCCs deficient in either GroP or Glc-GroP, whereas the sidewalls carry the fully mature SCC species decorated with Glc-GroP. Importantly, carbohydrate analysis of SCCs extracted from *S. mutans* WT, Δ*sccH* and Δ*sccH*:p*sccH* cell walls by mild acid hydrolysis established that deletion of *sccH* has no significant effect on total SCC content (Supplementary Fig. 9b). This fact excludes the possibility that the mislocalized binding of AtlA along the entire surface of Δ*sccH* bacteria is due to increased expression of SCC within the cell wall. To further test our hypothesis that *S. mutans* peptidoglycan is decorated with the SCC variants lacking either GroP or Glc-GroP, we separated the extracted SCCs using anion-exchange chromatography. The majority of WT SCC is negatively charged and binds tightly to the anion-exchange column, eluting as a broad peak (Fig. 4g). However, a substantial portion (∼15-20 %) is neutral, with very low phosphate content, and does not bind to the anion-exchange column. Our previous work on the GAC revealed similar results ^28^, suggesting that streptococcal species produce different forms of the cell wall polysaccharides. As we previously reported, the phosphate content in GAC and SCC is an indication of the presence of the GroP modification in the polysaccharide ^28^. Analysis of the glycosyl composition in the two fractions revealed that the column-bound fraction contains more Glc relative to Rha than the unbound fraction (Fig. 4h). As expected, SCC extracted from Δ*sccH* elutes as a single peak of neutral SCC, lacking detectable phosphate (Supplementary figure 12).

To further characterize the heterogeneity of the SCC variants, we examined the electrophoretic mobility of ANDS-conjugated polysaccharides extracted from WT, Δ*sccH*, Δ*sccN*, Δ*sccP* and Δ*sccN*Δ*sccP* (Fig. 4i). Since ANDS introduces a negative charge to SCC, this fluorescent tag allows examination of electrophoretic mobility of neutral polysaccharides extracted from the GroP-deficient mutants: Δ*sccH*, Δ*sccN* and Δ*sccN*Δ*sccP.* We observed the distinctive “laddering” of bands in the SCCs extracted from WT and Δ*sccP*, indicating the high level of heterogeneity of the polysaccharides. As expected and in agreement with the anion-exchange chromatography analysis, the SCCs extracted from Δ*sccH*, Δ*sccN* and Δ*sccN*Δ*sccP* migrated as a single band. Altogether, these data indicate that *S. mutans* WT produces SCC variants with different degrees of GroP modification.

### Z-ring positioning is controlled by GroP modification

The morphological abnormalities of Δ*sccH* and Δ*sccN* suggest a role for GroP modification of SCC in regulating either the assembly or correct positioning of the Z-ring. To test this hypothesis, we expressed FtsZ fused with tagRFP (FtsZ-tagRFP) from its chromosomal locus as the only copy in WT and Δ*sccH* cells (the *ftsZ-tagRFP* and Δ*sccH ftsZ-tagRFP* strains). To identify the localization of the GroP-deficient SCCs on the surface of *S. mutans*, we added AtlA-GFP exogenously to the *ftsZ-tagRFP* and Δ*sccH ftsZ-tagRFP* strains. As expected, we observed both FtsZ-tagRFP and AtlA-GFP at the mid-cell in newborn WT cells (Fig. 5a, stage 1). In the cells beginning to elongate, the AtlA-GFP signal splits into two parallel bands that move away from the division site where a single FtsZ-ring is present (Fig. 5a, stages 2 and 3). During middle-to-late cell division stage, a part of FtsZ splits into two rings and migrates to the mid-cell regions of the newly forming daughter cells, resulting in a distinctive three-band pattern (Fig. 5a, stage 4). AtlA-GFP was already localized at these sites, suggesting that the wall regions carrying the GroP-deficient SCCs are present at the equators of daughter cells prior to the arrival of FtsZ (Fig. 5a, stage 4). This localization pattern is consistent with our idea that the equatorial rings contain immature SCCs that lack GroP. Interestingly, during the late cell division stage, the AtlA-GFP signal was detected at the constricting septum (Fig. 5a, stage 5), indicating that the GroP-deficient SCCs are inserted in the septal cell wall during the pole maturation.

**Fig. 5.**
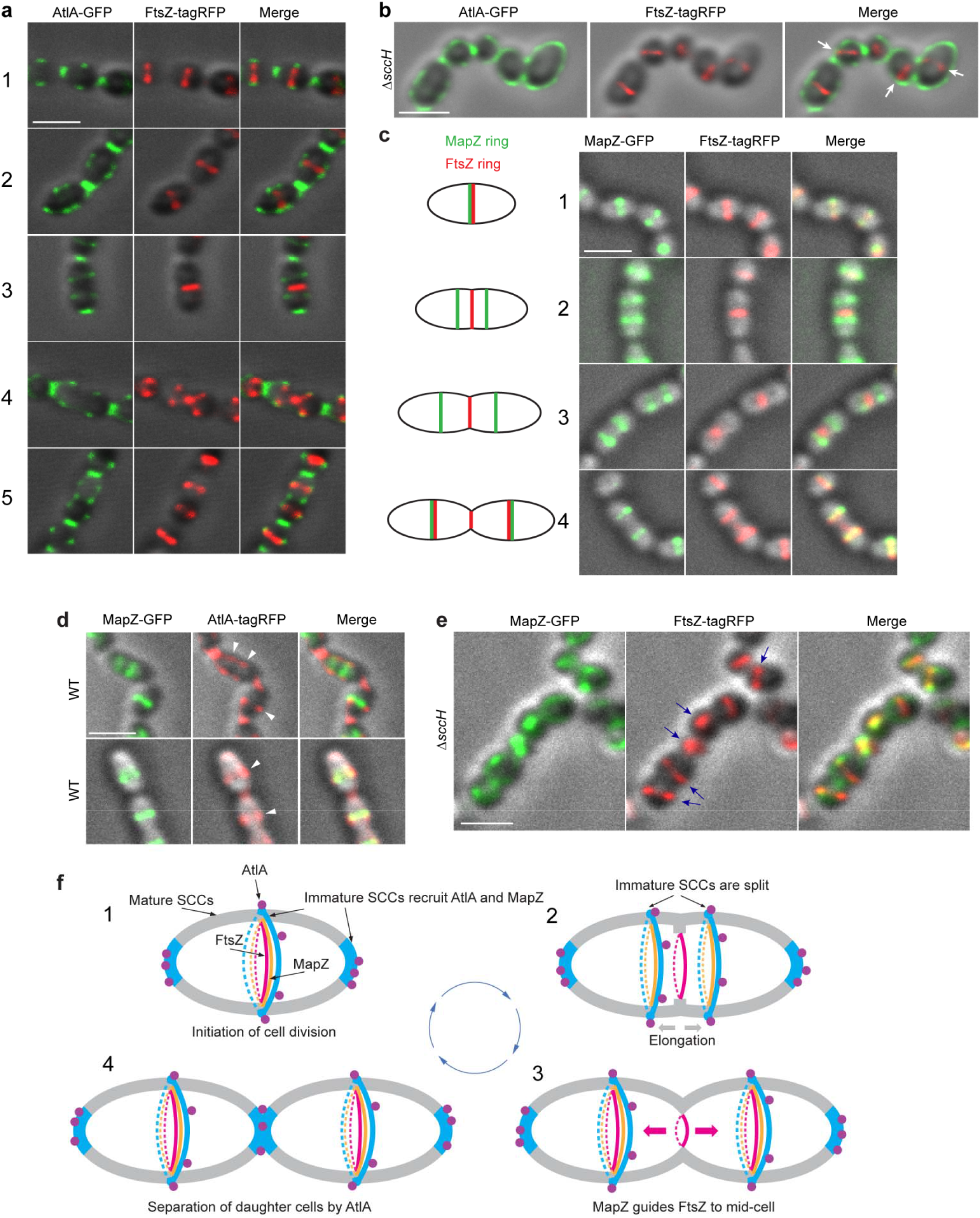
GroP modification of SCC controls the positioning of FtsZ- and MapZ-rings. (**a**) Localization of FtsZ in WT (strain *ftsZ-tagRFP*) cells at different stages of the cell cycle (designated 1 to 5). Images depict cells expressing FtsZ-tagRFP (red) labeled by AtlA-GFP protein (green). (**b**) Localization of FtsZ in Δ*sccH* (strain Δ*sccH ftsZ-tagRFP*) cells. Cells expressing FtsZ-tagRFP (middle panel, red) labeled with AtlA-GFP protein (left panel, green) were analyzed. An overlay of DIC, GFP, and tagRFP signals is shown in the far-right panel (merge). White arrows indicate mislocalized Z-rings. (**c**) Localization of MapZ (left panels, green) and FtsZ (middle panels, red) in WT (strain *ftsZ-tagRFP mapZ-GFP*) cells at different stages of cell cycle designated 1 to 4). WT cells expressing MapZ-GFP and FtsZ-tagRFP were analyzed. Schematic pictures (left panel) illustrate MapZ- and FtsZ-ring positions during different stages of the cell cycle. (**d**) Localization of MapZ (left panels, green) and SCCs labeled with AtlA-tagRFP protein (middle panels, red) in WT (strain *mapZ-GFP*) cells. White arrowheads depict the positions of equatorial SCCs labeled with AtlA-tagRFP. (**e**) Localization of MapZ (left panel, green) and FtsZ (middle panel, red) in Δ*sccH* (strain Δ*sccH ftsZ-tagRFP mapZ-GFP*) cells. Blue arrows indicate mislocalized Z-rings. Images are overlays of DIC and fluorescence images in **a, b, c, d** and **e**. An overlay of DIC, GFP, and tagRFP signals is shown in right panels in **a, c, d** and **e**. Scale bar is 1 µm in **a, b, c, d** and **e**. Representative images from at least three independent experiments are shown in **a, b, c, d** and **e**. (**f**) A schematic model of cell division in *S. mutans*. **1**. Cell division is initiated at mid-cell by recruitment and alignment of the FtsZ-ring with the equatorial ring. This process is guided by the MapZ-ring, which is recruited to mid-cell by immature SCCs present in the equatorial ring. AtlA binds to immature SCCs enriched in the cell poles and equatorial ring. **2**. Cell division machinery assembled at mid-cell synthesizes the first part of the septal wall. Concomitantly with this process, the equatorial ring is split in the middle presumably under turgor pressure and due to AtlA action. Synthesis of the lateral walls decorated with mature SCCs begins resulting in migration of new equatorial rings apart. The MapZ-rings follow the equatorial ring. **3**. The equatorial rings approach the equator of new daughter cells. MapZ guides FtsZ recruitment to new equatorial rings. **4**. Cell division machinery synthesizes the polar septal wall decorated with immature SCCs. AtlA is recruited to newly forming poles to separate the daughter cells. Cell walls decorated with mature and immature SCCs are shown in gray and blue, respectively. Purple circles indicate AtlA. FtsZ- and MapZ-rings are shown in red and orange colors, respectively.

In the Δ*sccH ftsZ-tagRFP* cells, the AtlA-GFP signal was evenly distributed along the cell surface, reflecting the presence of the GroP-deficient SCCs throughout the whole cell wall. While FtsZ was able to assemble into the Z-ring structures in the mutant cells, some Z-rings were often displaced from the cell center or not perpendicular to the cell’s axis (Fig. 5b and Supplementary figure 13a). These data indicate that the morphological defects of Δ*sccH* arise from a misplacement of the Z-ring.

### GroP modification regulates MapZ-ring positioning

The above results suggest that the correct positioning of Z-ring at the mid-cell depends on the presence of the GroP-deficient SCCs in the equatorial ring. Intriguingly, the cell shape alterations, together with the Z-ring misplacement observed in Δ*sccH*, are reminiscent of *S. pneumoniae* and *S. mutans* mutants lacking MapZ ^7-10^, implicating the GroP-deficient SCCs in the recruitment of MapZ to the equatorial ring. To test this hypothesis, we expressed the fusion of MapZ with GFP (MapZ-GFP) in the WT and Δ*sccH* genetic backgrounds resulting in the *mapZ-GFP* and Δ*sccH mapZ-*GFP strains. To analyze the correlation between MapZ and FtsZ, we generated the MapZ-GFP/FtsZ-tagRFP double expressing strains both in the WT and Δ*sccH* genetic backgrounds resulting in the *ftsZ-tagRFP mapZ-GFP* and Δ*sccH ftsZ-tagRFP mapZ* strains. The localization of the GroP-deficient SCCs on the surface of the *mapZ-GFP* strain was monitored by labeling bacteria with AtlA-tagRFP protein (the tagRFP fusion with the N-terminal Bsp repeats domain of AtlA), which was added exogenously. As expected from previous studies ^6-10^, in the WT newborn cells, the MapZ-ring coincides with the FtsZ-ring at mid-cell (Fig. 5c, stage 1). As cell division progresses, MapZ migrates as a pair of rings, parallel to the respective equators of newly forming daughter cells, while a single FtsZ-ring remains at the mid-cell of the parental cell (Fig. 5c, stages 2 and 3). At a later cell division stage, the majority of the FtsZ arrives at the future division sites in the daughter cells, where it co-localizes with MapZ (Fig. 5c, stage 4). AtlA-tagRFP associated with the WT cells with patterns very similar to that observed with AtlA-GFP being enriched at the poles and mid-cell regions (Fig. 5d). Importantly, the MapZ signal co-localizes with GroP-deficient SCCs indicated by AtlA-tagRFP labeling at mid-cell regions in newborn and elongating WT cells (Fig. 5d), which is consistent with the idea that these immature SCCs are present in the equatorial rings.

Strikingly, in the Δ*sccH* (strains Δ*sccH mapZ-GFP* and Δ*sccH ftsZ-tagRFP mapZ-GFP*) cells, MapZ did not assemble into the characteristic ring-like structures, but instead the MapZ-GFP fluorescent signal was dispersed throughout the membrane of the mutants (Fig. 5e and Supplementary figure 13b). As expected from the proposed function of MapZ in guiding FtsZ positioning ^7-10^, Z-rings were mislocalized in the Δ*sccH* cells (Fig. 5e). These results indicate that the enrichment of the GroP-deficient SCCs at the equatorial rings drives the recruitment and assembly of MapZ into the ring-like structures. The underlying mechanism likely relies on an exclusion strategy whereby decoration of SCC with GroP at distinct cellular positions provides the molecular signal to exclude recruitment of MapZ and FtsZ to these sites.

## Discussion

Despite years of intensive research, it remains unclear how oval-shaped Gram-positive bacteria generate equally sized daughter cells after division. This work uncovers a mechanism of molecular exclusion to position the cell division machinery in the human pathogen *S. mutans*. Our results are consistent with a model in which GroP-deficient SCCs are enriched at the equatorial rings and poles. Such controlled distribution of the polysaccharides provides the molecular cues for the simultaneous recruitment of cell division machinery as well as proper daughter cell separation.

Rha-containing cell wall polysaccharides are expressed by a wide variety of streptococcal, lactococcal and enterococcal species ^29^. We recently reported the discovery of GroP modification in the *S. mutans* and GAS cell wall polysaccharides, SCC and GAC, respectively ^28^. This modification is likely present in other streptococci since homologs of *sccH*, which encodes the *S. mutans* GroP transferase, is highly conserved among streptococcal species except for the *Streptococcus mitis* group ^28^. Structural studies of the cell wall polysaccharides isolated from streptococcal species, including *S. mutans*, proposed that each disaccharide repeating unit of the polyrhamnose backbone is modified with glycosyl side-chains ^29^. Previously, we have demonstrated that in the GAS, the GlcNAc side-chains of GAC are decorated with GroP ^28^. Here, we report that the side-chains of SCC are similarly decorated by GroP in *S. mutans*. Moreover, we now demonstrate that the structural architecture of Rha-containing cell wall polysaccharides has important functional implications for regulation of cell division in streptococci.

We observe that the GroP- and Glc side-chain-deficient mutants of *S. mutans* display severely impaired cell division, as represented by a large fraction of unequally sized cells. The morphological changes are accompanied by reduced viability and enhanced susceptibility to autolysis. These intriguing phenotypes prompted a search for responsible cell division proteins and autolysins. Cell wall pulldown studies, together with co-sedimentation analysis using SCC-depleted cell walls, identified SCC as the only detectable cell wall binding receptor for AtlA. This autolysin is involved in cell separation of daughter cells after cell division, autolysis, bacterial competence, and biofilm formation in *S. mutans* ^31-34,45^. Fluorescence anisotropy experiments conclusively demonstrate that AtlA-GFP binds to Glc- and GroP-modified SCCs, but the binding affinity for these variants is significantly lower than for those lacking the decorations. The analytical ultracentrifugation studies further revealed that all analyzed SCC variants, including the WT SCCs, interact with AtlA to form higher-order complexes. However, modification of SCC with either GroP or the Glc side-chains reduces the extent of the protein oligomerization. We show that the AtlA N-terminal domain which is composed of Bsp repeats, targets specifically the polyrhamnose backbone of SCC. This finding allowed us to employ a fusion of the AtlA N-terminal domain with a fluorescent protein as a tool to map the topological arrangements of different SCC species within the bacterial cell wall.

Using immunofluorescent microscopy we demonstrate that AtlA and AtlA-GFP are evenly distributed along the cell surface of the mutants lacking GroP and Glc side-chains, indicating that peptidoglycan is decorated with SCCs throughout the whole cell surface of *S. mutans*. Importantly, the absence of binding of AtlA-GFP to the Δ*rgpG* cells eliminates the possibility that AtlA recognizes additional structures on the bacterial surface. Finally, expression of the SCC polyrhamnose backbone is sufficient to confer AtlA binding, as demonstrated by expression in the non-natural host *E. coli*. The autolysin was found to be responsible for enhanced susceptibility to autolysis of the GroP- and Glc side-chain-deficient mutants. In newborn WT cells, AtlA is specifically targeted to cell poles, where the enzyme hydrolyzes peptidoglycan to allow daughter cell separation, and to cell equators. These observations lead to the conclusion that the mislocalization of AtlA and its strong binding to the cell wall causes the enhanced cell lysis of the mutants.

During the elongation phase, the SCC species labeled with the fluorescent AtlA fusion move from mid-cell of the parental cell, where the Z-ring is localized, toward the cell equators of the newly forming daughter cells. Subsequently, these SCCs arrive at the new division site before FtsZ. The apparent movement of the AtlA-labeled SCCs is the result of the newly synthesized cell wall being added at the septum, pushing the zone with these specific SCCs away from the cell division site. Furthermore, the AtlA-labeled SCCs co-localize with MapZ during the division cycle, indicating the presence of the distinct form of SCC in the equatorial rings. Characterization of SCCs extracted from *S. mutans* cell wall revealed a high degree of heterogeneity with a significant portion of SCC containing low or no GroP. These results, combined with the observed difference in AtlA localization in WT versus GroP-deficient bacteria, are consistent with the idea that peptidoglycan in the equatorial rings and cell poles is decorated with newly synthesized SCCs lacking GroP modifications. Note, the results do not exclude the possibility that these SCCs are also deficient for the Glc side-chain.

The highly organized spatial distribution of the GroP-deficient SCC variants further implies that the synthesis of the equatorial rings and cell poles requires a specific modulation of the divisome and/or elongasome complexes to produce cell wall decorated with immature SCCs at the early and final stages of cell division. In line with this idea, our co-localization analysis revealed that the translocation of FtsZ from the parental septum to the equators of the daughter cells coincides with the re-appearance of the AtlA-GFP-labeled SCCs at the parental septum, suggesting that the GroP-deficient cell wall material is incorporated into the septal wall at the last step of septum closure. This observation is in agreement with the time-lapse ultrastructural reconstruction of dividing *E. faecalis* indicating that the synthesis of septum occurs in two separate events — early in the division and after the elongation phase ^1^.

In ovococci, the peptidoglycan architecture at the equatorial rings has been proposed to direct the positioning of the cell division machinery through MapZ ^7,8^. Our analysis of MapZ localization in the GroP-deficient mutant revealed its impaired alignment with the equatorial rings. Interestingly, a similar localization defect was observed for MapZ lacking its extracellular domain in *S. pneumoniae* ^7^. Furthermore, we found that the delocalization of MapZ in the mutant cells causes a mis-orientation of the Z-rings, indicating that the cell shape deformations observed in the GroP-deficient mutants arise from a defect in septum placement. These findings highlight the pivotal role of GroP modification of SCC in cell division, leading to a model wherein immature SCCs, serve as landmarks for the recruitment of MapZ to equatorial rings and AtlA to equatorial rings and cell poles (Fig. 5f). Thus, this simple mechanism unifies septum positioning with its subsequent splitting. The importance of AtlA binding to equatorial rings is unclear at the moment. Early in the cell division cycle, AtlA might participate in the shaping of the new equatorial rings of the daughter cells by splitting the parental equatorial ring (Fig. 5f). A previous report, using electron microscopy imaging, has identified the equatorial rings as the zones of autolysin activity in *E. faecalis* ^46^. However, since the deletion of AtlA does not affect *S. mutans* cell shape, other mechanisms besides AtlA might be responsible for the generation of the daughter equatorial rings.

How MapZ recognizes cell wall decorated with immature SCCs is currently unknown but under investigation. Since MapZ homologs are also present in the species of the *S. mitis* group that do not express Rha-containing cell wall polysaccharides ^7,8^, it suggests that, in contrast to AtlA, MapZ does not interact directly with the cell wall polysaccharides. Note, that the morphological phenotype of GroP-deficient *S. mutans* does not depend on the specific glycosyl side chain to which GroP is attached since the mutant phenotype is complemented by the plasmid-based expression of the GAC-specific side-chain. This observation suggests that a major function of GroP in streptococcal cell wall is to provide a negative charge. Phosphate groups in WTA have been proposed to affect the packing density and rigidity of the cell wall due to electrostatic repulsion between the polysaccharides ^47^. Thus, it is possible that a cue for recognition of the equatorial ring by MapZ is an alteration in peptidoglycan density.

The Rha-containing cell wall polysaccharides are the functional homologs of canonical WTAs in streptococci because, similar to WTAs, they play significant roles in metal ion homeostasis, antimicrobial resistance ^28^, cell division and autolysis. The exact molecular mechanism by which WTAs regulate cell division and autolysis is not known. Much like GroP modification, WTAs are required for proper localization of autolysins by restricting the binding of autolysins from the entire bacterial surface and directing them to the specific cell wall sites where they are enzymatically active ^13,15,17^. Considering the similar function of WTAs and GroP modification of SCC, it is plausible that the here-described mechanism of the positioning of cell division proteins is widespread in Gram-positive bacteria. However, cell separation autolysins and the proteins bridging the Z-ring to cell wall likely differ between species.

## Methods

### Bacterial strains, growth conditions and media

All plasmids, strains and primers used in this study are listed in Supplementary Tables 1 and 2. Streptococci and *E. faecalis* were grown in BD Bacto Todd-Hewitt broth supplemented with 1% yeast extract (THY) without aeration at 37°C. *S. mutans* strains were grown either on THY or brain heart infusion (BHI) agar at 37°C with 5% CO_2_. *E. coli* strains were grown in Lysogeny Broth (LB) medium or on LB agar plates at 37°C. When required, antibiotics were included at the following concentrations: ampicillin at 100 μg mL^-1^ for *E. coli*; erythromycin at 5 μg mL^-1^ for *S. mutans*; chloramphenicol at 10 µg mL^-1^ for *E. coli* and 2 µg mL^-1^ for *S. mutans*; spectinomycin at 500 μg mL^-1^ for *S. mutans*; kanamycin at 300 μg mL^-1^ for *S. mutans*.

### Construction of mutant strains

To delete *sccN, S. mutans* Xc chromosomal DNA was used as a template for the amplification of two DNA fragments using two primers pairs: sccNup-BglII-f/sccNup-SalI-r and sccNdown-BamHI-f/sccNdown-XhoI-r (Supplementary Table 2). The first PCR product was digested with BglII/SalI and ligated into BglII/SalI-digested pUC19BXspec ^30^. The resultant plasmid was digested with BamHI/XhoI and used for ligation with the second PCR product that was digested with BamHI/XhoI. The resultant plasmid, pUC19BXspec-sccN, was digested with BglII and XhoI to obtain a DNA fragment containing the nonpolar spectinomycin resistance cassette flanked with the *sccN* upstream and downstream regions. The DNA fragment was transformed into *S. mutans* Xc by electroporation. The mutants were isolated as described below. The plasmids for the deletion of *sccP* and *atlA* were constructed using the same strategy described for *sccN* deletion with primer pairs listed in Supplementary Table 2.

To delete *rgpG* and *smaA* in the WT strain, *smaA* in the Δ*atlA* and Δ*sccN* backgrounds, *sccP* in the Δ*sccN* background, and *atlA* in the Δ*sccN* and Δ*sccH* backgrounds, we used a PCR overlapping mutagenesis approach, as previously described ^28^. Briefly, 600-700 bp fragments both upstream and downstream of the gene of interest were amplified with designed primers that contained 16-20 bp extensions complementary to the nonpolar antibiotic resistance cassette (Supplementary Table 2). The nonpolar spectinomycin, kanamycin, and erythromycin resistance cassettes were PCR-amplified from pLR16T, pOSKAR, and pHY304 (Supplementary Table 1), respectively. The two fragments of the gene of interest and the fragment with the antibiotic resistance cassette were purified using the QIAquick PCR purification kit (Qiagen) and fused by Gibson Assembly (SGA-DNA). A PCR was then performed on the Gibson Assembly sample using primers listed in Supplementary Table 2 to amplify the fused fragments.

To construct *S. mutans* strains expressing FtsZ fused at its C-terminus with a monomeric red fluorescent protein (tagRFP) ^48^, we replaced *ftsZ* with *ftsZ-tagRFP* at its native chromosome locus. A nonpolar kanamycin resistance cassette was inserted downstream of *ftsZ-tagRFP* to allow the selection of recombinant bacteria. The fragment encoding tagRFP was PCR-amplified from pTagRFP-N (Evrogen). The fragments of *ftsZ, tagRFP*, kanamycin resistance cassette, and the *ftsZ* downstream region were PCR-amplified using primers listed in Supplementary Table 2, purified and assembled as described above. *S. mutans* strains expressing MapZ fused at its N-terminus with a superfolder green fluorescent protein (GFP) ^36^ were constructed similarly, except that a nonpolar spectinomycin resistance cassette followed by the GAS *mapZ* promoter was inserted upstream of the *mapZ* fusion to allow the *mapZ* expression and selection of recombinant bacteria. The fragments of the *mapZ* upstream region, spectinomycin resistance cassette, GFP, and *mapZ* were PCR-amplified using primers listed in Supplementary Table 2. The assembled PCR fragment was transformed into corresponding *S. mutans* strains by electroporation. The transformants were selected either on THY or BHI agar containing the corresponding antibiotic. Double-crossover recombination was confirmed by PCR and Sanger sequencing using the primers listed in Supplementary Table 2.

### Complementation of ΔsccN

To complement Δ*sccN, sccN* was amplified from *S. mutans* Xc chromosomal DNA using primer pair sccN-HindIII-f/sccN-BglII-r, digested using HindIII/BglII, and ligated into HindIII/BglII-digested pDC123, yielding p*sccN*. To replace the SCC side-chain with the GAC side-chain, a part of the GAC operon required for the addition of the GlcNAc side-chains and GroP ^30^ was expressed on pDC123 in Δ*sccN*. GAS 5448 genomic DNA was used to amplify the *gacHIJKL* region using primer pairs A109-f/A101-r (Supplementary Table 2). The PCR fragment was digested using XhoI/BamHI, and ligated into XhoI/BamHI-digested pDC123, yielding p*gacHIJKL*. All plasmids were confirmed by sequencing analysis (Eurofins MWG Operon) before electroporation into *S. mutans*. We found that one *E. coli* clone carrying p*gacHIJKL* acquired a frameshift resulting in a stop codon at leucine 49 in GacI. The plasmid was designated p*gacHI*JKL* and used as a negative control in our experiments. Plasmids were transformed into Δ*sccN* by electroporation. Chloramphenicol resistant single colonies were picked and checked for the presence of p*sccN* or p*gacHIJKL* by PCR, yielding strains Δ*sccN:*p*sccN*, Δ*sccN*:p*gacHIJKL* and Δ*sccN*:p*gacHI*JKL.*

### Construction of the plasmids for expression of AtlA-GFP, AtlA-cGFP, AtlA-tagRFP and GFP

To create a vector for expression of the AtlA N-terminal Bsp domains fused with a superfolder GFP (AtlA-GFP fusion protein), two overlapping amplicons were generated. The first fragment encoding the N-terminal Bsp domains was amplified from *S. mutans* Xc chromosomal DNA using the primer pair atlA_fus-F and atlA_fus-R. The second fragment encoding GFP was amplified from pHR-scFv-GCN4-sfGFP-GB1-NLS-dWPRE (Supplementary Table 1) using the primer pair gfp_fus-F and gfp_fus-R. The PCR products were ligated into NcoI/HindIII-digested pRSF-NT vector (Supplementary Table 1) using Gibson assembly, resulting in pKV1527 for expression of AtlA-GFP.

To create a vector for expression of the AtlA N-terminal Bsp domains fused with colorless GFP (AtlA-cGFP fusion protein), two PCR products were amplified using primer pairs atlA_fus-F/Y66L_R and Y66L_F/gfp_fus-R, and pKV1527 as a template to introduce Y66L mutation into GFP ^49^.

To create a vector for expression of the AtlA N-terminal Bsp domains fused with tagRFP (AtlA-tagRFP fusion protein), two PCR products were amplified using primer pairs atlA_fus-F/atl_tRFP_R and tRFP_fusF/tRFP_fusR using *S. mutans* Xc chromosomal DNA and pTagRFP-N, respectively, as the templates. The corresponding PCR products were ligated into NcoI/HindIII-digested pRSF-NT vector using Gibson assembly resulting in pKV1556 and pKV1572.

To create a vector for expression of GFP, a PCR product was amplified using primer pair sfGFP_BspH/gfp_fus-R and pKV1527 as a template. The PCR product was digested with BspHI/HindIII and ligated into the NcoI/HindIII-digested pRSF-NT vector resulting in pKV1532.

### Protein expression and purification

To purify AtlA-GFP, AtlA-cGFP, AtlA-tagRFP and GFP, *E. coli* Rosetta (DE3) cells carrying the respective plasmids were grown in LB at 37°C to OD_600_ = 0.4-0.6 and induced with 0.25 mM isopropyl β-D-1-thiogalactopyranoside (IPTG) at 27°C for approximately 4 hours. Bacteria were lysed in 20 mM Tris-HCl pH 7.5, 300 mM NaCl by a microfluidizer cell disrupter. The proteins were purified by Ni-NTA chromatography followed by size exclusion chromatography (SEC) on a Superdex 200 column in 20 mM HEPES pH 7.5, 100 mM NaCl.

### Viability assay

Exponentially growing bacteria (OD_600_ of 0.5) were pushed ten times through a 26G 3/8 syringe to break bacterial clumps. Bacteria were serially diluted in phosphate-buffered saline (PBS) and plated on THY agar for enumeration.

### Triton X-100-induced autolysis assay

Overnight cultures of *S. mutans* were diluted (1:100) into fresh THY broth and grown to an OD_600_ of 0.5. The autolysis assay was primarily performed, as outlined in ^50^. Cells were allowed to autolyze in PBS containing 0.2% Triton X-100. The autolysis was monitored after 2, 4, and 21 hours as a decrease in OD_600_. Results were normalized to the OD_600_ at time zero (OD_600_ of 0.5).

### Isolation of cell walls and sacculi

The cell walls were isolated from exponential phase cultures by the SDS-boiling procedure and lyophilized, as previously described ^30^. The sacculi were obtained using the same protocol except that the bead beating step was omitted. The cell wall and sacculi were free of lipoteichoic acid (LTA), proteins, lipids, and nucleic acids.

### Binding of AtlA-GFP to intact bacteria and cell wall material

To study AtlA-GFP binding to intact bacteria, 10 mL of the overnight-grown bacteria were washed twice with PBS, resuspended in 1 mL of PBS, and incubated with 0.1 mg mL^-1^ AtlA-GFP for 1 hour with agitation. As a control, GFP of the same concentration was used in parallel with each experiment. Sample aliquots were assayed to determine the total fluorescence. Then samples were centrifuged (16,000 g, 3 min), and 100 µL of the supernatant was assayed for fluorescence. To determine the fluorescence of the pellet, the supernatant fluorescence was subtracted from the total fluorescence of the sample. Controls without bacterial cells indicated that the sedimentation of AtlA-GFP under these conditions was negligible. Data are presented as a percentage of fluorescence of the pellet normalized to the total fluorescence of the sample.

To examine AtlA-GFP binding to purified cell walls, 0.5 mg of lyophilized cell wall was incubated with 0.1 mg mL^-1^ AtlA-GFP in 0.5 mL of PBS. The experiment was conducted as described above.

### Pulldown of cell wall-associated proteins

*S. mutans* (1 L) were grown to an OD_600_ of 1.0, collected by centrifugation (5,000 g, 10 min), washed three times with PBS, and resuspended in 25 mL of urea solution (8 M urea, 20 mM Tris-HCl pH 7.5, 150 mM NaCl). The sample was rotated at room temperature for 1 hour, and then centrifuged (3,200 g, 10 min). The supernatant was dialyzed overnight at 4°C to remove urea and centrifuged again (3,200 g, 10 min). The supernatant was transferred to a fresh centrifuge tube and incubated with 10 mg of lyophilized cell wall with rotation for two hours. The cell wall was collected by centrifugation (3,200 g, 10 min) and washed three times with PBS. The proteins, retained in the cell wall, were dissolved in 0.5 mL of SDS sample buffer, and separated on 10% SDS-PAGE. Protein identification was performed at the Proteomics Core Facility (University of Kentucky) by liquid chromatography with tandem mass spectrometry (LC-MS/MS) analysis using an LTQ-Orbitrap mass spectrometer (Thermo Fisher Scientific) coupled with an Eksigent Nanoflex cHiPLC system (Eksigent) through a nanoelectrospray ionization source. The LC-MS/MS data were subjected to database searches for protein identification using Proteome Discoverer software V. 1.3 (Thermo Fisher Scientific) with a local MASCOT search engine.

### Release of SCCs from cell wall by mild acid hydrolysis

SCC was released from purified cell walls by mild acid hydrolysis following N-acetylation, as previously described for WTA of *Lactobacillus plantarum* ^51^ with some modifications. N-acetylation was conducted with lyophilized cell wall (4 mg) and 2 % acetic anhydride in 1 mL of saturated NaHCO_3_. After incubation at room temperature overnight, the reactions were diluted with 2 vol water, and sedimented at 50,000 g, 10 min. The cell walls were further washed by resuspension with 3 mL water followed by sedimentation at 50,000 g, 10 min, three times. N-acetylated cell walls were resuspended with 0.02 N HCl (0.2 mL), and heated to 100°C for 20 min. The reactions were cooled on ice, neutralized by the addition of 4 µL 1 N NaOH, and sedimented at 50,000 g, 10 min. The supernatant fraction was removed and reserved. The pellet was resuspended in 0.2 mL water and re-sedimented. The supernatant fractions were combined and either analyzed or purified further by a combination of SEC and ion-exchange chromatography.

### Fractionation of SCCs on DEAE-Toyopearl

SCCs were released from purified cells walls by mild acid hydrolysis and fractionated on BioGel P150 equilibrated in 0.2 N sodium acetate, pH 3.7, 0.15 M NaCl, as previously described using BioGel P10 ^28^. The fractions containing SCCs were combined and concentrated by centrifugation over an Amicon Ultra - 15 Centrifugal Filter (KUltracel -3K). The retentate was desalted by repeated cycles of dilution with water and centrifugation. The concentrated SCCs (∼0.5 mL) were applied to a 1×18 cm column of TOYOPEARL DEAE-650M (TOSOH Bioscience), equilibrated in 10 mM Tris-Cl, pH 7.4, and fractions of 2 mL were collected. After 20 fractions, the column was eluted with an 80 mL gradient of NaCl (0-0.5 M). Appropriate aliquots were analyzed for Rha and Glc by anthrone assay.

### Derivatization of SCCs with 7-amino-1,3-naphthalenedisulfonic acid (ANDS)

SCCs purified by a combination of SEC and ion-exchange chromatography were fluorescently tagged at the reducing end by reductive amination with ANDS as previously described ^52^. Reaction mixtures contained 30-100 nmol of SCCs (lyophilized), 0.75 mM ANDS and 0.5 M NaCNBH_4_ in 0.05 mL acetic acid/DMSO at 7.5/50 (%). Following overnight incubation at 37°C, the reactions were desalted by centrifugation on an Amicon Ultra Centrifugal Filter (3,000 NMWL). Derivatized SCCs were further purified by SEC over a Superdex 75 10/300 GL column (GE Healthcare Bio-Sciences AB).

### Glycerol and phosphate assays

The total phosphate content of SCCs was determined by the malachite green method following digestion with perchloric acid, as previously described ^28^. To determine glycerol, SCCs were incubated with 2 N HCl at 100°C for 2 hours. The samples were neutralized with NaOH in the presence of 62.5 mM HEPES pH 7.5. Glycerol concentration was measured using the Glycerol Colorimetric assay kit (Cayman Chemical) according to the manufacturer’s protocol.

### Modified anthrone assay

Total Rha and Glc contents were estimated using a minor modification of the anthrone procedure. Reactions contained 0.08 mL of aqueous sample and water and 0.32 mL anthrone reagent (0.2 % anthrone in concentrated H_2_SO_4_). The samples were heated to 100°C, 10 min, cooled in water (room temperature), and the absorbance at 580 nm and 700 nm was recorded. The chromophore produced from Rha has a relatively discreet absorbance maximum and is essentially zero at 700 nm. The absorbance of the Glc chromophore at 700 nm is approximately equal to its absorbance at 580 nm. Therefore the contribution of Glc to the absorbance at 580 nm can be estimated from its absorbance at 700 nm and subtracted from the absorbance at 580 nm to obtain the Rha-specific signal. Rha and Glc concentrations were estimated using L-Rha and D-Glc standard curves, respectively.

### Glycosyl composition analysis

Glycosyl composition analysis of SCC samples was performed at the Complex Carbohydrate Research Center (University of Georgia, Athens, GA) by combined gas chromatography/mass spectrometry (GC-MS) of the per-O-trimethylsilyl derivatives of O-methyl glycosides of the monosaccharides produced by acidic methanolysis as described previously ^30^. Likewise, a similar assay was performed following in-house pretreatment with HF, as described below. Appropriate aliquots were supplemented with 2 nmol inositol (internal standard), taken to dryness, and incubated with 25 µL 51% HF, 4°C, 16 hours. Following HF treatment, the samples were evaporated under a stream of air (the evaporating HF was captured by bubbling through a trap containing 1 M KOH), dissolved in water, transferred to screw-cap tubes equipped with Teflon lined caps and dried again. 0.2 mL 1 N HCl in methanol (formed by the drop-wise addition of acetyl chloride into rapidly-stirring anhydrous methanol) was added, the tubes were tightly sealed and incubated at 80°C, 16 hours. Following methanolysis, the reactions were neutralized with ∼3-5 mg AgCO_3_, centrifuged, and the supernatant transferred to a fresh tube. To re-N-acetylate amino sugars, the samples were taken to dryness under a stream of air, evaporated out of 0.5 mL of methanol, and re-dissolved in 0.1 mL of methanol containing 10 % pyridine and 10 % acetic anhydride. The reactions were dried, trimethylsilylated with 25 μL Tri-Sil TH (Sigma Aldrich), and analyzed by methane chemical ionization GC-MS using a Thermo ISQ mass spectrometer interfaced with a gas chromatograph, equipped with a 15 m Equity 1701 glass capillary column and helium carrier gas.

### GlcNAc analysis

GlcNAc content was assayed using the Megazyme Glucosamine Assay Kit according to the manufacturer’s instructions with some minor modifications. Partially purified polysaccharide (∼40-60 nmol Rha) was hydrolyzed in 40 µL 2 N HCl, 100°C, 2 hours, neutralized with 10 N NaOH (to ∼pH 7.0 by pH paper) and adjusted to a final volume of 50 µL with water. The acid hydrolysis de-acetylates GlcNAc to generate glucosamine. An aliquot (5 µL) of the neutralized hydrolysate was diluted with 171 µL water and mixed with a total of 24 µL of Megazyme Glucosamine Assay Kit Enzyme Mix. Absorbance at 340 nm was recorded after incubation at room temperature for 40 min. Glucosamine content was determined by comparison with a glucosamine standard curve and verified by comparison with GlcNAc standards treated similarly as the polysaccharide.

### FACS analysis

AtlA-GFP (5 µL, 3.8 mg mL^-1^) or GFP (5 µL, 3.5 mg mL^-1^) were added to 100 µL of *E. coli* CS2775 ^41^ or PHD136 [*E. coli* CS2775 harboring pRGP1 plasmid ^41^] (OD_600_ = 0.4). After 20 minutes of incubation on ice, the cells were centrifuged at 20,800 g, 5 min. The pellet was washed twice with PBS, resuspended in PBS, fixed with paraformaldehyde (4% final concentration), 4°C, 20 min, and then washed once with PBS. The cells were resuspended in 0.3% BSA in PBS and immediately analyzed by flow cytometry (BD LSRFortessa). Anti-GAC antibodies conjugated with FITC (ABIN238144, antibodies-online, titer 1:50) were used as a positive control to confirm polyrhamnose expression in *E. coli* strain PHD136. The FACS data were analyzed using flowJo version 10.

### Scanning electron microscopy (SEM)

Exponentially growing bacteria (OD_600_ of 0.7) were harvested by centrifugation (3,200 g, 10 min), washed once with 20 mM HEPES, 0.5% BSA buffer (pH 8.0), resuspended in PBS, fixed with paraformaldehyde (4% final concentration), 4°C, 15 min, and then pipetted onto microscope slide cover glasses coated with poly-L-lysine. Following one hour incubation, the cover glasses were washed 3 times with PBS. Bacteria were dehydrated stepwise in a gradient series of ethanol (35%, 50%, 70%, and 96% for 20 min each and then 100% overnight), followed by critical point drying with liquid CO_2_ in a Leica EM CPD300. Samples were coated with about 5 nm of platinum controlled by a film-thickness monitor. SEM images were performed in the immersion mode of an FEI Helios Nanolab 660 dual beam system.

### Fluorescent and differential interference contrast (DIC) microscopy

To determine the bacterial regions targeted by AtlA-GFP and AtlA-tagRFP, we utilized fluorescent microscopy. Exponentially growing bacteria (OD_600_ of 0.7) were fixed with paraformaldehyde (4% final concentration), 4°C, 15 min, pipetted onto microscope slide cover glasses coated with poly-L-lysine, and allowed to settle for one hour at room temperature. Bacteria were incubated with AtlA-GFP or AtlA-tagRFP (20 µg mL^-1^) for 15 min at room temperature. As a control, GFP of the same concentration was used in parallel with each experiment. The samples were washed four times with PBS, dried at room temperature, and mounted on a microscope slide with ProLong Diamond Antifade Kit with DAPI (Invitrogen). Samples were imaged on a Zeiss LSM 880 using Airyscan mode and a Leica SP8 equipped with 100X, 1.44 N.A. objective, DIC optics, and “lightning” post-processing. Images were processed with Airyscan processing and the “lightning” processing tool, respectively. Samples with *S. mutans* cells expressing MapZ-GFP and FtsZ-tagRFP were prepared similarly. They were imaged on a Leica SP8. Images were deconvolved using Huygens Professional software.

Immunofluorescent microscopy was used to monitor AtlA on the cell surface of *S. mutans*. Exponentially growing bacteria (OD_600_ of 0.8) were fixed with 2.5% (v/v) paraformaldehyde, 0.03% glutaraldehyde in 30 mM phosphate buffer (pH 8.0), transferred onto poly-L-lysine coated cover glasses and incubated at room temperature for one hour. The samples were washed three times with PBS and incubated in buffer containing 20 mM Tris-HCl pH 7.5, 10 mM EDTA, 50 mM Glc, and lysozyme (0.1 mg mL^-1^) for 30 min. After washing twice with PBS, the samples were air-dried, and dipped in cold methanol, -20°C, 5 min. The samples were blocked with 2% (w/v) bovine serum albumin in PBS (BSA-PBS), room temperature, 2 hours, and incubated with polyclonal anti-AtlA antibodies ^33^ (1:600) in BSA-PBS, 4°C, overnight. The samples were then washed 15 min five times with PBS+0.1% Tween-20 (PBST) buffer and incubated with Alexa Fluor 488-conjugated goat anti-rabbit antibodies (1:100) in BSA-PBS, room temperature, 2 hours. After extensive washing with PBST buffer, the specimens were air-dried and assembled on microscope slides mounted with ProLong Diamond Antifade Kit with DAPI. Micrographs were taken on a Leica SP8 confocal microscope.

To determine the length and width of cells, exponentially growing bacteria (OD_600_ of 0.7) were imaged on a Leica SP8 confocal microscope. ImageJ software was used to measure the sizes of cells.

### Fluorescence anisotropy

Reactions (150 µL) containing 0.5 µM ANDS-labeled polysaccharide and 0-10 µM AtlA-cGFP were incubated at room temperature for 30 minutes in 20 mM Tris 7.5, 300 mM NaCl. The anisotropy was then measured on a Fluoromax-4 (Horiba) photon-counting steady-state fluorometer at 25°C using an excitation wavelength of 310 nm and the following emission at 450 nm with slit widths of 5 nm and an integration time of 1 second. All measurements are an average of three independent replicates. Curves terminate at 10 µM of AtlA-cGFP due to aggregation issues at higher protein concentration. The single-site total binding equation in Graphpad Prism 8 was used to fit the binding of the polysaccharide isolated from the Δ*sccN*Δ*sccP* mutant. Lower-bound estimates of the K_d_ for the other three SCC species were determined by fitting to the binding equation after fixing the maximal binding limit, B_max_, to that of Δ*sccN*Δ*sccP.*

### Analytical ultracentrifugation

Sedimentation velocity (SV) experiments were performed in a Beckman ProteomeLab XL-I at 20°C using absorbance optics at 278 nm in an An-60Ti rotor at 32,000 rpm until complete sedimentation of sample occurred. The analysis was conducted using Sedfit 16.1c ^53,54^ using the c(s) data model and expressed using sedimentation coefficient distributions. Each fit rmsd was 0.005 or lower. GUSSI 1.4.1 ^55^ was used for data visualization. SV data were also fitted using Wide Distribution Analysis (WDA) in SedAnal v7.11 ^56^. WDA distributions were computed from 6.10 cm to 7.00 cm with an increment of 0.01 cm. The radial range plotted was 6.40 - 6.60 cm in 0.01 cm increments, with a 2% smoothing algorithm applied (equivalent to 4 scans). The weight-average sedimentation coefficient (sw) was computed using a range of 2-100 S. Partial specific volume (v-bar) was 0.725 ml g^-1^, and the solution density (ρ) was 0.998 g mL^-1^.

### Statistical analysis

Unless otherwise indicated, statistical analysis was carried out on pooled data from at least three independent biological repeats. Statistical analysis of data was performed using one-way ANOVA, 2-way ANOVA, and two-tailed Student’s t-test as described for individual experiments. A *P*-value equal to or less than 0.05 was considered statistically significant.

### Data availability

All data generated during this study are included in the article and supplementary information files or will be available from the corresponding author upon reasonable request.

## Acknowledgments

The authors thank Dr. Sang-Joon Ahn (University of Florida) for the kind gift of anti-AtlA antibodies, Dr. John F. Timoney (University of Kentucky) and Dr. Johannes Huebner (von Hauner Children’s Hospital, LMU) for providing *S. equi* and *E. faecalis*, respectively, Dr. Jeffrey M. Bosken and Dr. Edward D. Hall (University of Kentucky) for the use of the Thermo Scientific GC-MS instrument and Dr. Catalina Velez-Ortega (University of Kentucky) for the access to Leica SP8 confocal microscope. This work was supported by NIH grants R56 AI135021 from the NIAID and R01 DE028916 from the NIDCR (to NK), R01 GM094363 from the NIGMS (to ABH) and R01 DC014658 from the NIDCD (to GIF), Tenovus Scotland Large Research Grant T17/17 and University of Dundee Wellcome Fund 105606/Z/14/Z (to SAC and HCD), Wellcome and Royal Society Grant 109357/Z/15/Z (to HCD). Scanning electron microscopy was performed at the Electron Microscopy Center, which belongs to the National Science Foundation NNCI Kentucky Multiscale Manufacturing and Nano Integration Node, supported by ECCS-1542174. Carbohydrate composition analysis at the Complex Carbohydrate Research Center was supported by the Chemical Sciences, Geosciences and Biosciences Division, Office of Basic Energy Sciences, U.S. Department of Energy grant (DE-FG02-93ER20097) to Parastoo Azadi. The funders had no role in study design, data collection and interpretation, or the decision to submit the work for publication.

## Author contributions

SZ, CTC, JSR, SAC, AEY, ABH, NMvS, HCD, GIF, KVK, and NK designed the experiments. SZ, CTC, JSR, SAC, AEY, HCD, KVK, and NK performed functional and biochemical experiments. SZ and GIF performed microscopy analysis. NK, KVK, and NMvS constructed plasmids and isolated mutants. SZ, CTC, JSR, SAC, AEY, ABH, HCD, KVK, and NK analyzed the data. NK wrote the manuscript with contributions from all authors. All authors reviewed the results and approved the final version of the manuscript.

## Competing interests

The authors declare no competing interests.

## Notes

### Competing Interest Statement

The authors have declared no competing interest.

