## Supplementary Information for "Modification of cell wall polysaccharide spatially controls cell division in *Streptococcus mutans*"

**Supplementary Table 1.** Bacterial strains and plasmids.

| Strain or plasmid | Description | Reference |
|---|---|---|
| ***Bacteria*** | | |
| GAS | 5448 GAS, M1T1-serotype strain | 1 |
| *S. equi* | *S. equi* subsp. *equi* CF32 was isolated from a submandibular abscess of a horse | 2 |
| *S. agalactiae* | GBS A909, a serotype Ia, sequence type 7 GBS strain widely used in laboratory investigations | ATCC |
| *E. faecalis* | V583, vancomycin-resistant clinical isolate | ATCC |
| ***Streptococcus mutans*** | | |
| Xc | Serotype *c* strain, wild-type (WT) | 3 |
| Δ*sccH* | *sccH* deletion mutant (has a nonpolar erythromycin resistance cassette inserted in *sccH*), $Ery^R$ | 4 |
| Δ*sccH*:p*sccH* | Δ*sccH* is complemented with p*sccH* carrying WT *sccH*, $Ery^R$, $Cat^R$ | 4 |
| Δ*sccN* | *sccN* deletion mutant (has a nonpolar spectinomycin resistance cassette inserted in *sccN*), $Spec^R$ | This study |
| Δ*sccN*:p*sccN* | Δ*sccN* is complemented with p*sccN* carrying WT *sccN*, $Spec^R$, $Cat^R$ | This study |
| Δ*sccP* | *sccP* deletion mutant (has a nonpolar spectinomycin resistance cassette inserted in *sccP*), $Spec^R$ | This study |
| Δ*sccN*Δ*sccP* | *sccN sccP* double-gene deletion mutant (has a nonpolar spectinomycin and erythromycin resistance cassettes inserted in *sccN* and *sccP*, respectively), $Spec^R$ $Ery^R$ | This study |
| Δ*atlA* | *atlA* deletion mutant (has a nonpolar spectinomycin resistance cassette inserted in *atlA*), $Spec^R$ | This study |
| Δ*sccH*Δ*atlA* | *sccH atlA* double-gene deletion mutant (has a nonpolar erythromycin and spectinomycin resistance cassettes inserted in *sccH* and *atlA*, respectively), $Spec^R$ $Ery^R$ | This study |
| Δ*sccN*Δ*atlA* | *sccN atlA* double-gene deletion mutant (has a nonpolar spectinomycin and kanamycin resistance cassettes inserted in *sccN* and *atlA*, respectively), $Spec^R$ $Kan^R$ | This study |
| Δ*smaA* | *smaA* deletion mutant (has a nonpolar kanamycin resistance cassette inserted in *smaA*), $Kan^R$ | This study |
| Δ*sccN*Δ*smaA* | *sccN smaA* double-gene deletion mutant (has a nonpolar spectinomycin and kanamycin resistance cassettes inserted in *sccN* and *smaA*, respectively), $Spec^R$ $Kan^R$ | This study |
| Δ*smaA*Δ*atlA* | *smaA atlA* double-gene deletion mutant (has a nonpolar kanamycin and spectinomycin resistance cassettes inserted in *smaA* and *atlA*, respectively), $Spec^R$ $Kan^R$ | This study |
| Δ*rgpG* | *rgpG* deletion mutant (has a nonpolar spectinomycin resistance cassette inserted in *rgpG*), $Spec^R$ | This study |
| *ftsZ-tagRFP* | *ftsZ* was replaced with *ftsZ-tagRFP* followed by kanamycin resistance cassette in the WT background, $Kan^R$ | This study |

| | | |
|---|---|---|
| Δ*sccH ftsZ-tagRFP* | *ftsZ* was replaced with *ftsZ-tagRFP* followed by kanamycin resistance cassette in the Δ*sccH* background, $Ery^R$, $Kan^R$ | This study |
| *mapZ-GFP* | *mapZ* was replaced with *mapZ-GFP* (MapZ fused at the N-terminus with GFP) in the WT background. A nonpolar spectinomycin resistance cassette inserted upstream of *mapZ-GFP.* $Spec^R$ | This study |
| Δ*sccH mapZ-GFP* | *mapZ* was replaced with *mapZ-GFP* (MapZ fused at the N-terminus with GFP) in the Δ*sccH* background. A nonpolar spectinomycin resistance cassette inserted upstream of *mapZ-GFP.* $Ery^R$, $Spec^R$ | This study |
| *mapZ-tagRFP* | *mapZ* was replaced with *mapZ-tagRFP* (MapZ fused at the N-terminus with tagRFP) in the WT background. A nonpolar spectinomycin resistance cassette inserted upstream of *mapZ-tagRFP.* $Spec^R$ | This study |
| Δ*sccH mapZ-tagRFP* | *mapZ* was replaced with *mapZ-tagRFP* (MapZ fused at the N-terminus with tagRFP) in the Δ*sccH* background. A nonpolar spectinomycin resistance cassette inserted upstream of *mapZ-tagRFP.* $Ery^R$, $Spec^R$ | This study |
| *ftsZ-tagRFP mapZ-GFP* | *mapZ* was replaced with *mapZ-GFP* (MapZ fused at the N-terminus with GFP) in the *ftsZ-tagRFP* genetic background. A nonpolar spectinomycin resistance cassette inserted upstream of *mapZ-GFP.* $Kan^R$, $Spec^R$ | This study |
| Δ*sccH ftsZ-tagRFP mapZ-GFP* | *mapZ* was replaced with *mapZ-GFP* (MapZ fused at the N-terminus with GFP) in the Δ*sccH ftsZ-tagRFP* genetic background. A nonpolar spectinomycin resistance cassette inserted upstream of *mapZ-GFP.* $Ery^R$, $Kan^R$, $Spec^R$ | This study |
| Δ*sccN*:p*gacHIJKL* | Δ*sccN* is complemented with p*gacHIJKL*, $Spec^R$, $Cat^R$ | This study |
| Δ*sccN*:p*gacHI*JKL* | Δ*sccN* is complemented with p*gacHI*JKL* $Spec^R$, $Cat^R$ | This study |
| ***Escherichia coli*** | | |
| DH5α | *E. coli* cells used for cloning | Invitrogen |
| Rosetta (DE3) | *E. coli* cells used for protein expression | Novagen |
| CS2775 | *rfaS2007::Tnlac* derivative of CS2767; $Kan^R$ | [5] |
| PHD136 | CS2775 carrying pRGP1; $Kan^R$, $Ery^R$ | This study |
| ***Plasmids*** | | |
| pTagRFP-N | A mammalian expression vector encoding a monomeric red fluorescent protein (TagRFP) | Evrogen |
| pHR-scFv-GCN4-sfGFP-GB1-NLS-dWPRE | Expression vector encoding a superfolder green fluorescent protein (GFP) | Gift from Ron Vale (Addgene plasmid # 60906) [6] |
| pRSF-NT | A modified pRSF-Duet1 (Novagen) vector that allows the creation of N-terminus His-tagged proteins with a TEV protease cleavage site | [7] |
| pKV1527 | pRSF-NT derived plasmid expressing AtlA-GFP (N-terminal domain of AtlA fused with GFP at the C-terminus. | This study |

| | | |
|---|---|---|
| | AtlA is fused at the N-terminus with a His-tag followed by a TEV protease recognition site) | |
| pKV1556 | pRSF_AtlA-GFP derived plasmid expressing AltA fused with colorless GFP, AtlA-cGFP | This study |
| pKV1572 | pRSF-NT derived plasmid expressing AtlA-tagRFP (N-terminal domain of AtlA fused with tagRFP at the C-terminus. AtlA is fused at the N-terminus with a His-tag followed by a TEV protease recognition site) | This study |
| pKV1532 | pRSF-NT derived plasmid expressing GFP fused at the N-terminus with a His-tag followed by a TEV protease recognition site | This study |
| pDC123 | *E. coli-streptococcus* shuttle vector, JS-3 replicon, $Cat^R$. | [8] |
| p*sccH* | pDC123 derived plasmid expressing *sccH* | [4] |
| p*sccN* | pDC123 derived plasmid expressing *sccN* | This study |
| p*gacHIJKL* | pDC123 derived plasmid expressing *gacHIJKL* | This study |
| p*gacHI*JKL* | pDC123 derived plasmid expressing *gacHI*JKL* (with a stop codon at L49 in GacI) | This study |
| pUC19BXspec | Derivative of pUC19BX expressing *aadA* (spectinomycin resistance cassette) with own rbs | [9] |
| pUC19BXspec-sccN | Derivative of pUC19BXspec expressing *aadA* flanked with *sccN* 5' and 3' regions | This study |
| pUC19BXspec-sccP | Derivative of pUC19BXspec expressing *aadA* flanked with *sccP* 5' and 3' regions | This study |
| pUC19BXspec-atlA | Derivative of pUC19BXspec expressing *aadA* flanked with *atlA* 5' and 3' regions | This study |
| pRGP1 | Plasmid containing 10.0-kb BstEII fragment of *S. mutans* Xc47 chromosomal DNA which includes *sccA* through *sccG*, $Ery^R$ | [10] |
| pLR16T | Vector encoding spectinomycin resistance cassette | [11] |
| pOSKAR | Vector encoding kanamycin resistance cassette | [12] |
| pHY304 | Vector encoding erythromycin resistance cassette | [13] |

**Supplementary Table 2.** Primers

| Primer | Sequence[a,b] | Genetic manipulations |
|---|---|---|
| sccNup-BglII-f | GCGTAAGATCTGGTTCTGACAGTCGTCTCTC | *sccN* deletion with nonpolar spectinomycin resistance cassette |
| sccNup-SalI-r | CGCTGCGTCGACGGTTTCTTCCTCATTATAAC | |
| sccNdown-BamhI-f | GATTTACAGGATCCGCCAGAATTG | |
| sccNdown-XhoI-r | GCGCGCTCGAGGCAACAAAATTTAGAATCAACAAC | |
| sccPup-BglII-f | GCGTAAGATCTCGCTCTTTATTTGCGAAAAC | *sccP* deletion with nonpolar spectinomycin resistance cassette |
| sccPup-SalI-r | CGCTGCGTCGACCTCTTCATTGTAAGCTGGAAG | |
| sccPdown-BamhI-f | CGTCTGGATCCGCTCGTCAAGCTGGTGCGATTG | |
| sccPdown-XhoI-r | GCGCGCTCGAGCCTAACGCAAAAGACAAGGC | |
| atlA-BglII-f | GCGTAAGATCTGCACCTAAAAATCTGGATAAG | *atlA* deletion with nonpolar spectinomycin resistance cassette |
| atlA-SalI-r | CGCTGCGTCGACGGGTCAATGCTAATGGAATC | |
| atlA-BamhI-f | CGTCTGGATCCGCAGCAAACACAGGAACAG | |
| atlA-XhoI-r | GCGCGCTCGAGCATTTAATATCATCTTGACC | |
| RgpG-f | GAGTCTGACGCTTATCACATG | *rgpG* deletion with nonpolar spectinomycin resistance cassette |
| Spec-RgpG-r1 | **CACTATTTTGGTCGAC**CAGCATTAGTAATGGCAACAAC | |
| RgpG-Spec-f1 | GCCATTACTAATGCTG**GTCGACCAAAATAGTGAGGAGG** | |
| Spec-RgpG-f2 | **AAAATTATAAGGATCC**GATGATTAGTCTGACCACTATG | |
| RgpG-Spec-r2 | GTCAGACTAATCATC**GGATCCTTATAATTTTTTAATCTG** | |
| RgpG-r | CAAGTACAGACATTGTACGCTC | |
| Ery-sccP-r1 | **TAATTTAACTTCAATTCC**CTCTTCATTGTAAGCTGGAAG | *sccP* deletion with nonpolar erythromycin resistance cassette |
| sccP-Ery-f1 | GCTTACAATGAAGAG**GGAATTGAAGTTAAATTAGATG** | |
| Ery-sccP-f2 | **GAGGAAATAATTCTATG**GCTCGTCAAGCTGGTGCGATTG | |
| sccP-Ery-r2 | CACCAGCTTGACGAGC**CATAGAATTATTTCCTCCCG** | |
| smaA-f | GGCATTGAGCAATTGGTGCAAG | *smaA* deletion with nonpolar kanamycin resistance cassette |
| Kan-smaA-r1 | **CAGTATTTAAAGATACC**GCTCCAAATGCATATTTGCG | |
| smaA-Kan-f1 | AAATATGCATTTGGAGC**GGTATCTTTAAATACTGTAG** | |
| Kan-smaA-f2 | **TGAATTGTTTTAGTAC**CTGCTTTAAATGTCGATGAC | |

| | | |
|---|---|---|
| smaA-Kan-r2 | CGACATTTAAAGCAG**GTACTAAAACAATTCATCCAG** | |
| smaA-r | CCAATAACAACATAACGACGG | |
| Kan-AtlA-r1 | **CAGTATTTAAAGATACC**GTCAATGCTAATGGAATC | *atlA* deletion with nonpolar kanamycin resistance cassette |
| AtlA-Kan-f1 | TCCATTAGCATTGAC**GGTATCTTTAAATACTGTAG** | |
| Kan-AtlA-f2 | T**GAATTGTTTTAGTAC**GCAGCAAACACAGGAACAG | |
| AtlA-Kan-r2 | TTCCTGTGTTTGCTGC**GTACTAAAACAATTCATCCAG** | |
| sccN-check-f | GTTATAATGAGGAAGAAACC | Verification of Δ*sccN* |
| sccN-check-r | CAATTCTGGCGGATCCTG | |
| sccPcheck-f | CTTCCAGCTTACAATGAAGAG | Verification of Δ*sccP* |
| sccPcheck-r | CAATCGCACCAGCTTGACGAGC | |
| atlA-check-f | GATTCCATTAGCATTGACCC | Verification of Δ*atlA* |
| atlA-check-r | CTGTTCCTGTGTTTGCTGC | |
| rgpG-check-f | GCTTGGTGCTGTTATTATTTG | Verification of Δ*rgpG* |
| rgpG-check-r | GAAACCAGCAATAGCAAAGATC | |
| smaA-check-f | GTATATCATAATGATAGGAGTG | Verification of Δ*smaA* |
| smaA-check-r | GTAGGGTTCACATCTTCTTTGG | |
| sccN-HindIII-f | CGCGCAAGCTTGAAAAGGTGAGATGGCAAGAAGG | Construction of p*sccN* plasmid |
| sccN-BglII-r | GCGTAAGATCTGCCTTTATCCTTTTTCCTTAAC | |
| A101-r | TACCTCGAGGTTTAATGATAATATCTAAAAATAGTACTC | Construction of p*gacHIJKL* plasmid |
| A109-f | TACGGATCCCACAAAACTCTATATTACATGCGATTGAG | |
| atlA_fus-F | AACCTTTACTTCCAGGGCGCCATGGATGAGCAAAATCAATCCTTAAG | Construction of pKV1527 and pKV1532 plasmids |
| atlA_fus-R | GCTACCGCTTCCAGATGGTAGAGCAACAGCAGG | |
| gfp_fus-F | GCTCTACCATCTGGAAGCGGTAGCAAAGGAG | |
| gfp_fus-R | TTAAGCATTATGCGGCCGCAAGCTTATTTGTAGAGCTCATCCATGCC | |
| sfGFP_BspH-f | GAGATCATGAGCAAAGGAGAAGAACTTTTCAC | |
| Y66L_F | CTTGTCACTACTCTGACCCTGGGTGTTCAATGCTTTTC | Y66L mutation in GFP, construction of pKV1556 |
| Y66L_R | GAAAAGCATTGAACACCCAGGGTCAGAGTAGTGACAAG | |

| | | |
|---|---|---|
| atl_tRFP_R | CGCCCTTAGACACGCTACCGCTACCAGATGGTAGAGCAAC | Construction of pKV1572 |
| tRFP_fusF | CATCTGGTAGCGGTAGCGTGTCTAAGGGCGAAGAGC | |
| tRFP_fusR | TTAAGCATTATGCGGCCGCAAGCTTAATTAAGTTTGTGCCCCAG | |
| FtsZ-f | GGAATCGGTATCGGTACTGG | *ftsZ* replacement with *ftsZ-tagRFP*. FtsZ is fused at the C-terminus with TagRFP using GSGS linker. |
| TagRFP-FtsZ-r1 | CACAGCGCTACCGCTTCCAGATGAACGATTCTTAAAGAAAGGAG | |
| FtsZ-TagRFP-f1 | CCTTTCTTTAAGAATCGTTCATCTGGAAGCGGTAGCGCTGTGTCTAAGGGCGAAGAGC | |
| TagRFP-kan-F2 | CTGGGGCACAAACTTAATTGA**GGTATCTTTAAATACTGTAG** | |
| Kan-TagRFP-R2 | **CAGTATTTAAAGATACC**TCAATTAAGTTTGTGCCCCAG | |
| Kan-FtsZ-F2 | **TGAATTGTTTTAGTAC**ATAATGGATTTACAAGCAAATAAAG | |
| FtsZ-kan-R2 | CTTGTAAATCCATTAT**GTACTAAAACAATTCATCCAG** | |
| FtsZ-r | GGTTGAGCCATTTCGAATAGC | |
| mapZ-F | CGATACAACAGGACCAAGTC | *mapZ* replacement with *mapZ-GFP*. MapZ is fused at the N-terminus with GFP using SGSGS linker |
| mapZ-spec-F1 | CGTGTGCTTAGATAG**GTCGACCAAAATAGTGAGGAG** | |
| spec-mapZ-R1 | **CTCACTATTTTGGTCGAC**CTATCTAAGCACACGTTTGACAC | |
| Spec-prGFP-F2 | TGTATCATTGATAATAATACTGAAATATAATTTTGGTAAAAAGATTGACCGTTCGGAGAGTAAAGAATGAGCAAAGGAGAAGAACTTTTC | |
| prGFP-spec-R2 | ATTATATTTCAGTATTATTATCAATGATACAGAAAAATAGTTCAAAAAGCAAGTATTTAAATACCAATA**CCTTATAATTTTTTTAATCTG** | |
| GFP-mapZ-F3 | CTCTACAAATCTGGAAGCGGTAGCTCAGAAAAGGAAAAAAATCC | |
| mapZ-GFP-R3 | GGATTTTTTTCCTTTTCTGAGCTACCGCTTCCAGATTTGTAGAG | |
| mapZ-r | GCTGTGTGATGGCTCAATTG | |

[a] Restriction sites are underlined.
[b] Extensions complementary to the antibiotic resistance cassettes, TagRFP, GFP are in bold, red and green, respectively

**Supplementary Table 3.** Cell size analysis[a]

| | WT | Δ*sccH* | Δ*sccH*:p*sccH* | Δ*sccN* | Δ*sccN*:p*sccN* | Δ*sccN*: p*gacHIJKL* | Δ*sccN*: p*gacHI*JKL* | Δ*atlA* | Δ*sccH* Δ*atlA* | Δ*smaA* |
|---|---|---|---|---|---|---|---|---|---|---|
| Number of cells | 85 | 94 | 136 | 132 | 113 | 133 | 82 | 196 | 107 | 158 |
| Average length | 0.88±0.11 | 0.74±0.14 | 1.07±0.16 | 0.78±0.18 | 1.00±0.15 | 0.89±0.11 | 0.77±0.13 | 0.87±0.12 | 0.81±0.17 | 0.92±0.14 |
| Average width | 0.59±0.06 | 0.60±0.06 | 0.64±0.05 | 0.68±0.07 | 0.63±0.05 | 0.64±0.07 | 0.65±0.07 | 0.59±0.05 | 0.67±0.08 | 0.65±0.05 |
| Average aspect ratio | 1.50±0.23 | 1.23±0.2 | 1.67±0.28 | 1.15±0.23 | 1.57±0.25 | 1.39±0.21 | 1.19±0.22 | 1.48±0.22 | 1.22±0.27 | 1.43±0.24 |
| % of cells with aspect ratio<=1 | 0 | 12 | 0 | 23 | 0 | 1 | 22 | 0 | 27 | 1 |
| Number of minicells | 2 | 20 | 0 | 19 | 0 | 1 | 17 | 0 | 18 | 1 |
| Minicells (%) | 2 | 21 | 0 | 14 | 0 | 1 | 21 | 0 | 17 | 1 |

[a] Cell size (μm) was measured by ImageJ software, and the results were analyzed by GraphPad Prism 3.0. Values are reported with standard deviation. Minicells represent all cells shorter than 0.646 μm (the minimal length of the WT cells).

**Supplementary Table 4**. Genes encoding the Bsp repeats proteins identified in the genomes of Firmicutes

| Bacteria | protein | Catalytic domain | Genomic location |
|---|---|---|---|
| *Streptococcus mutans* | AtlA | none | NAFC[1] |
| | SmaA | GH25 | NAFC |
| | LytF | CHAP | Downstream of SCC biosynthesis operon |
| *Streptococcus gallolyticus* UCN34 | GALLO_0625 | none | NAFC |
| | GALLO_1197 | peptidase C39 | NAFC |
| | GALLO_1368 | GH25 | Downstream of Rha polysaccharide biosynthesis operon |
| *Streptococcus agalactiae* 2603V/R | SAG1350 | none | NAFC |
| | SAG1206 | Peptidase C39 | NAFC |
| *Streptococcus gordonii* str. Challis substr. CH1 | SGO_2013 | Glyco_hydro_25 and CHAP | NAFC |
| | SGO_2094 | none | NAFC |
| *Streptococcus pasteurianus* ATCC 43144 | SGPB_1289 | GH25 | Downstream of Rha polysaccharide biosynthesis operon |
| | SGPB_1290 | none | |
| *Streptococcus sanguinis* SK36 | SSA_0036 | CHAP | In the middle of Pur operon |
| | SSA_0257 | Peptidase C39 | NAFC |
| | SSA_0860 | CHAP | NAFC |
| *Streptococcus parasanguinis* CC87K | HMPREF1195_01729 | CHAP | Downstream of Rha polysaccharide biosynthesis operon |
| | HMPREF1195_01730 | GH25 and Peptidase C39 | Downstream of Rha polysaccharide biosynthesis operon |
| | HMPREF1195_01895 | CHAP | In the middle of Pur operon |
| *Streptococcus suis* 05ZYH33 | SSU05_1294 | N-acetylmuramoyl-L-alanine amidase | In the middle of Rha polysaccharide biosynthesis cluster |
| *Streptococcus thermophilus* LMD-9 | STER_1030 | none | NAFC |
| | STER_0495 | none | NAFC |
| *Streptococcus uberis* 0140J | SUB0705 | N-acetylmuramoyl-L-alanine amidase | Downstream of Rha polysaccharide biosynthesis operon |
| *Streptococcus salivarius* M18 | SSALIVM18_m10337 | none | NAFC |
| | SSALIVM18_m10577 | CHAP and DUF4775 (domain of unknown function) | NAFC |
| | SSALIVM18_m10597 | VanY | NAFC |
| | SSALIVM18_02065 | none | NAFC |
| | SSALIVM18_02095 | VanY | NAFC |
| | SSALIVM18_04946 | GH25 | NAFC |

| | | | |
|---|---|---|---|
| | SSALIVM18_07441 | none | NAFC |
| *Streptococcus anginosus* C238 | SANR_RS09335 | VanY | NAFC |
| | SANR_RS09350 | CHAP | NAFC |
| | SANR_RS00260 | CHAP | In the middle of Pur operon |
| *Streptococcus criceti* HS-6 | STRCR_2363 | none | NAFC |
| | STRCR_1831 | VanY | NAFC |
| | STRCR_0770 | Peptidase C39 | NAFC |
| | STRCR_1202 | GH25 | NAFC |
| *Streptococcus porcinus* str. Jelinkova 176 | STRPO_0572 | N-acetylmuramoyl-L-alanine amidase | Downstream of Rha polysaccharide biosynthesis operon |
| | STRPO_0176 | CHAP | NAFC |
| *Streptococcus intermedius* B196 | SIR_RS14950 | Peptidase C39 and GH25 | NAFC |
| | SIR_RS10010 | CHAP | In the middle of Pur operon |
| *Streptococcus parauberis* NCFD 2020 | SPB_0281 | N-acetylmuramoyl-L-alanine amidase | Downstream of Rha polysaccharide biosynthesis operon |
| | SPB_1046 | Peptidase C39 | NAFC |
| | SPB_1655 | pfam13354 (beta-lactamase2) | NAFC |
| | SPB_0450 | none | NAFC |
| *Streptococcus lutetiensis* 033 | KE3_0988 | Peptidase C39 | NAFC |
| | KE3_1252 | GH25 | Downstream of Rha polysaccharide biosynthesis operon |
| | KE3_1253 | CHAP | Downstream of Rha polysaccharide biosynthesis operon |
| *Streptococcus urinalis* 2285-97 | STRUR_1425 | N-acetylmuramoyl-L-alanine amidase | Downstream of Rha polysaccharide biosynthesis operon |
| | STRUR_1426 | CHAP | Downstream of Rha polysaccharide biosynthesis operon |
| | STRUR_0608 | none | NAFC |
| | STRUR_1063 | CHAP | NAFC |
| | STRUR_1755 | DMP1 (domain of unknown function) | NAFC |
| *Enterococcus cecorum* DSM 20682 | I567_0095 | none | NAFC |
| | I567_02337 | none | NAFC |
| | I567_02446 | GH25 | In the middle of Rha polysaccharide biosynthesis cluster |
| | I567_02335 | Peptidase C39 | NAFC |
| | I567_01135 | PI-PLCc_GDPD_SF | NAFC |
| | I573_01472 | Peptidase C39 | NAFC |

| | | | |
|---|---|---|---|
| *Enterococcus sulfureus* ATCC 49903 | OMY_00864 | N-acetylmuramoyl-L-alanine amidase | Downstream of polysaccharide biosynthesis operon |
| *Enterococcus columbae DSM 7374* | OMW_00898 | NLPC_P60 | NAFC |
| | OMW_00899 | Peptidase C39 | NAFC |
| *Ruminococcus gnavus CAG:126* | BN481_01689 | Glucosaminidase | Downstream of Rha polysaccharide biosynthesis operon |
| | BN481_02281 | Transglutaminase-like | NAFC |
| *Roseburia hominis* | RHOM_12165 | N-acetylmuramoyl-L-alanine amidase and glucosaminidase | Downstream of polysaccharide biosynthesis operon |
| | RHOM_12160 | Transglutaminase-like | Downstream of polysaccharide biosynthesis operon |
| *Lactococcus reticulitermitis* | RsY01_1802 | N-acetylmuramoyl-L-alanine amidase and lysozyme subfamily 2 | Downstream of Rha polysaccharide biosynthesis operon |
| *Paenibacillus alvei DSM 29* | PAV_5c01530 | none | NAFC |
| | PAV_5c03180 | none | NAFC |
| *Acetobacterium woodii DSM 1030* | Awo_c02360 | N-acetylmuramoyl-L-alanine amidase and glucosaminidase | In the middle of Rha polysaccharide biosynthesis cluster |
| | Awo_c02370 | NlpD and peptidase M23 | Downstream of Rha polysaccharide biosynthesis operon |
| | Awo_c23170 | CHAP | NAFC |
| | Awo_c16100 | CHAP | NAFC |
| | Awo_c11470 | none | NAFC |

[1]NAFC — no apparent functional coupling

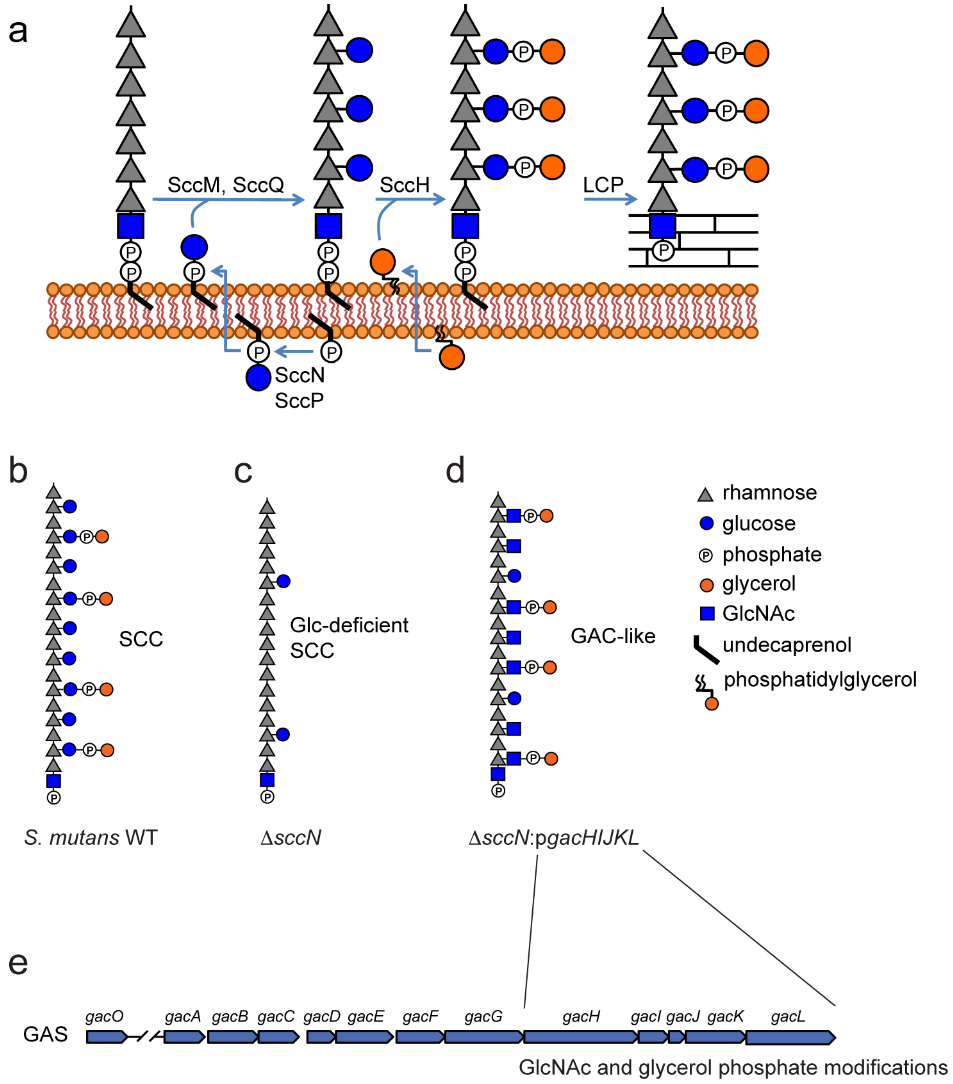

**Supplementary Fig. 1. Construction of *S. mutans* mutant expressing SCC with the GAC GlcNAc side-chains.**

(**a**) The proposed mechanism of SCC modification with the Glc side-chains and GroP. In the inner leaflet of the membrane, SccN and SccP produce Glc-phosphate-undecaprenol, which is then transported across the plasma membrane to the outer leaflet by an unknown flippase. Subsequently, SccM and SccQ most likely transfer Glc to polyrhamnose backbone using Glc-phosphate-undecaprenol as glycosyl donor. Protein members of the LytR-CpsA-Psr (LCP) phosphotransferase family presumably attach SCC to peptidoglycan. Lastly, SccH transfers GroP to the Glc side-chains. Several steps of this biosynthetic scheme are still speculative, such as the transfer of Glc by SccM and SccQ, and further research is required to confirm this hypothetical pathway definitively. However, the overall organization is consistent with the proposed and largely proven mechanism of GAC modification with the GlcNAc-GroP side-chains [9]. (**b**) The Glc side-chains of SCC are modified by GroP. (**c**) The Δ*sccN* mutant is GroP and Glc side-chain deficient. (**d**) Expression of *gacHIJKL* [a part of the GAC operon required for the addition of the GlcNAc side-chains and GroP [9]] in Δ*sccN* results in substitution of the SCC side-chains with the GAC side-chains. Phosphate groups in the SCC structures are involved in the phosphodiester bond linking glycerol to the glycosyl side-chain. (**e**) Schematic representation of the GAC biosynthetic gene cluster.

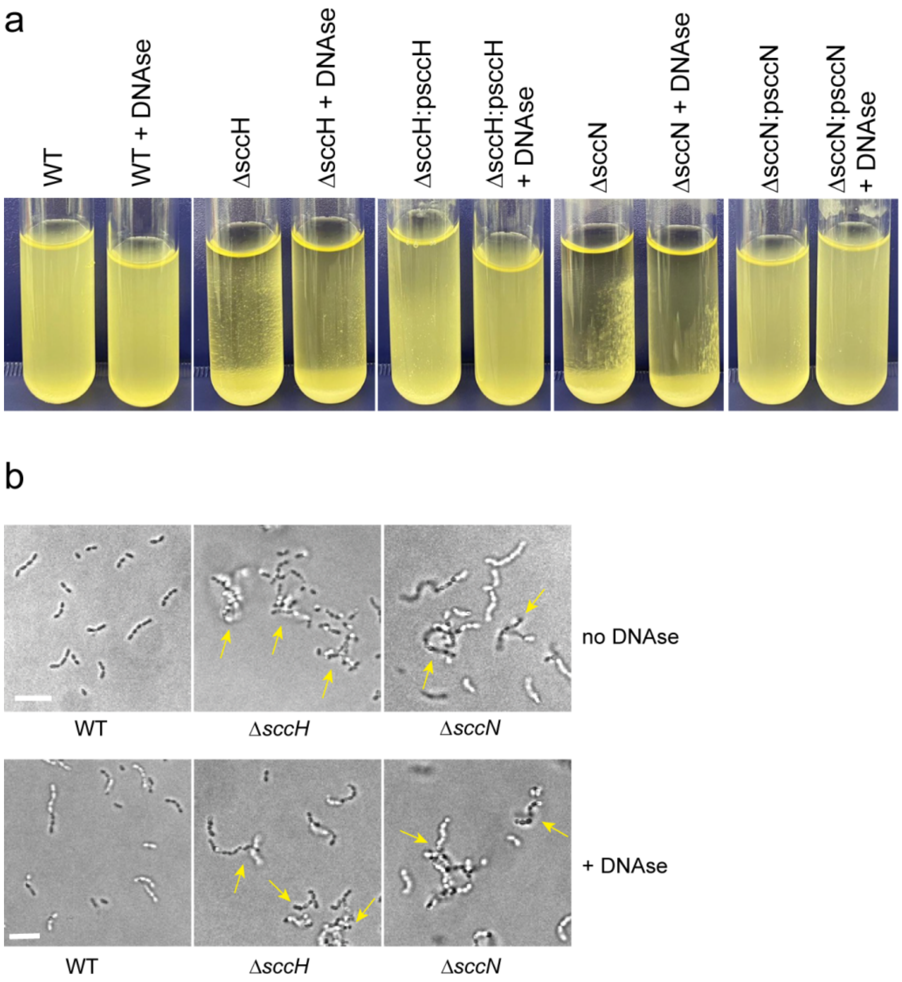

**Supplementary Fig. 2. Effect of DNAse on the self-aggregation of GroP-deficient mutants.** (**a**) Effect of DNAse on the growth phenotype of WT, Δ*sccH*, Δ*sccH*:p*sccH*, Δ*sccN* and Δ*sccH*:p*sccN*. It has been reported that the deletion of *sccN* resulted in lysis-independent DNA release in *S. mutans* [14,15], which caused bacterial self-aggregation that could be reversed after treatment of the mutant with DNAse [15]. To investigate the role of DNA in the self-aggregation of Δ*sccH* and Δ*sccN*, we grew bacteria overnight in THY broth in presence/absence of 5 μg $mL^{-1}$ DNAse. (**b**) Differential interference contrast (DIC) images of the WT, Δ*sccH*, and Δ*sccN* bacteria grown in THY broth overnight in presence/absence of 5 μg $mL^{-1}$ DNAse. Yellow arrows indicate bacterial aggregates. Top panels in **b** (cells with no DNAse) are the same as in Figure 2b. Scale bar is 5 μm. The experiments depicted in **a** and **b** were performed independently three times and yielded the same results. Representative image from one experiment is shown.

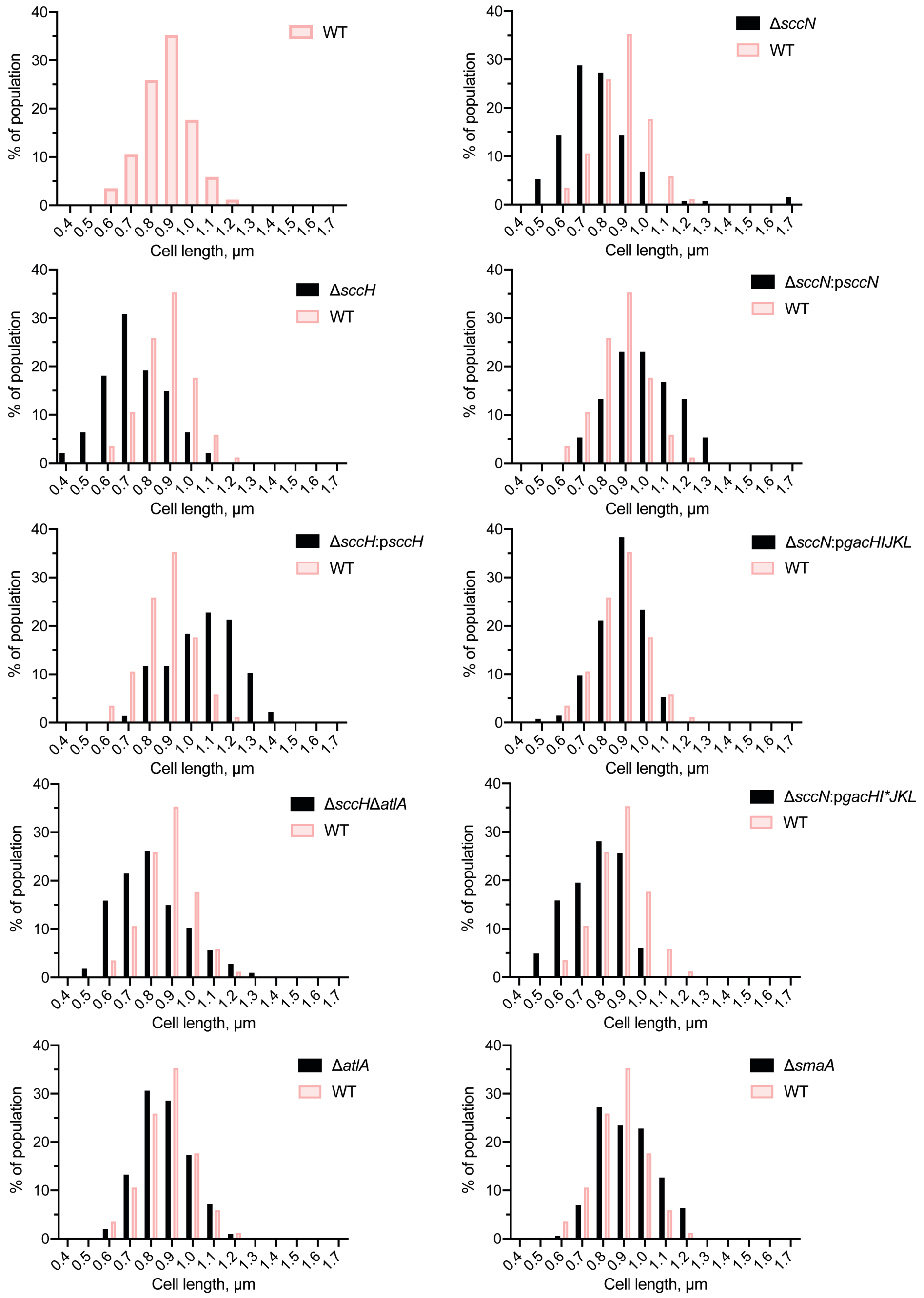

**Supplementary Fig. 3. Cell length distribution analysis**.

The histogram shows the distribution of cells of the WT (red) and the corresponding mutant (black) in distinct size classes. Numbers on the X axis indicate the medium size of the cell in the corresponding class. Cell size (µm) was measured by ImageJ software, and the results were analyzed by GraphPad Prism 3.0.

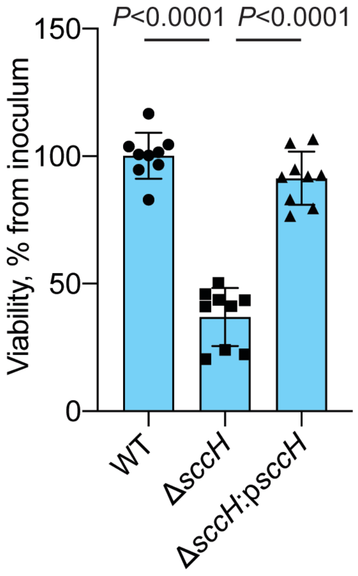

**Supplementary Fig. 4. SccH is important for cell viability of *S .mutans*.**

Viable colony count analysis. *S. mutans* WT, Δ*sccH* and Δ*sccH:psccH* were grown in THY to $OD_{600}$ = 0.5. Cells were pushed ten times through a 26G 3/8 syringe to break bacterial clumps. Bacteria were serially diluted in phosphate-buffered saline and plated on THY agar for enumeration. Results were normalized to the WT colony number. Columns and error bars represent the mean and S.D., respectively. n = 9 biologically independent replicates. *P*-values were calculated by one-way ANOVA with Tukey's multiple comparisons test.

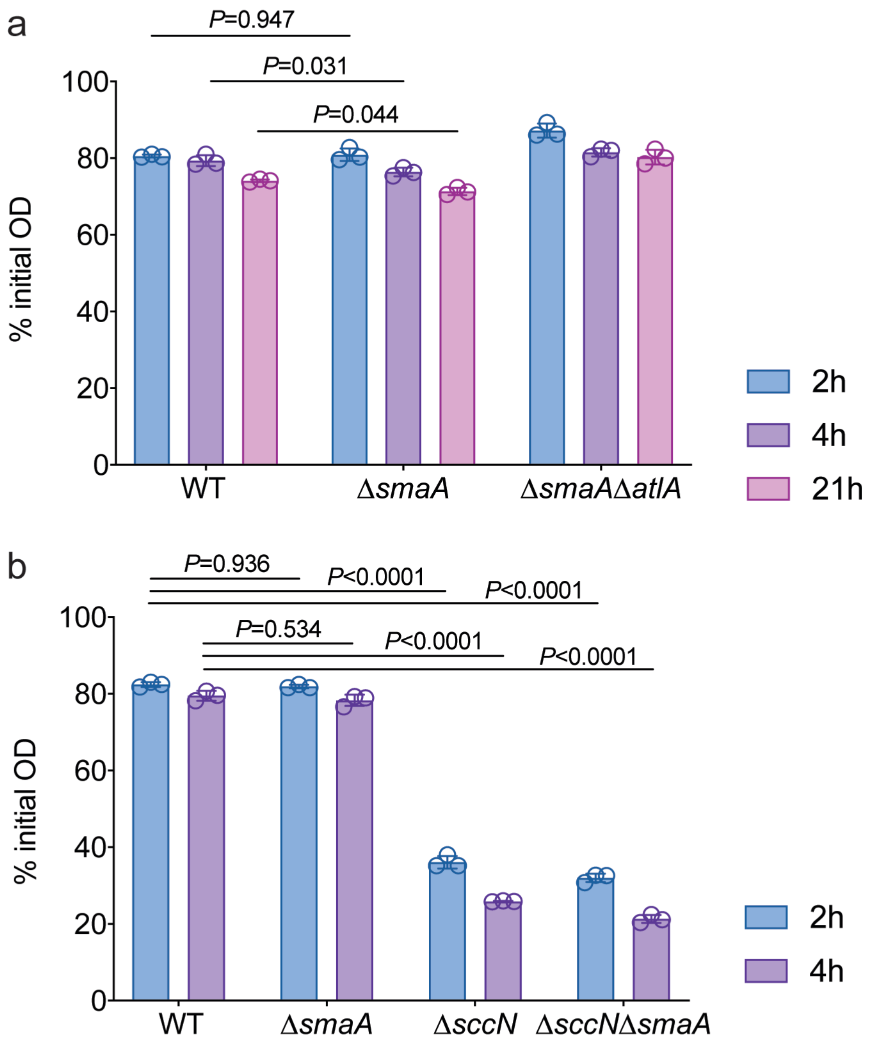

**Supplementary Fig. 5. SmaA is not involved in Triton X-100-induced autolysis of *S. mutans* WT and Δ*sccN***

The autolytic activity of *S. mutans* WT, Δ*smaA*, Δ*smaAΔatlA*, Δ*sccN* and Δ*sccNΔsmaA.* Exponentially growing strains were allowed to autolyze in 0.1% Triton X-100. The autolysis was monitored after 2, 4, and 21 h as the decrease in $OD_{600}$. Results were normalized to the $OD_{600}$ at time zero ($OD_{600}$ of 0.5). Columns and error bars represent the mean and S.D., respectively (n = 3 biologically independent replicates). *P*-values were determined by two-way ANOVA with Tukey's multiple comparisons test.

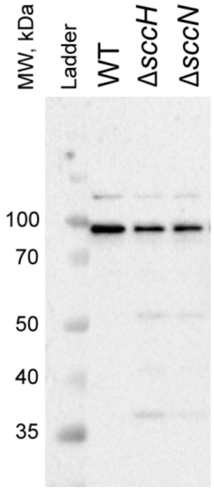

**Supplementary Fig. 6. SccH and SccN do not regulate the expression of AtlA.**

AtlA was extracted from the cell surface of the WT, Δ*sccH*, and Δ*sccN* bacteria with 4% SDS as outlined in [16]. Proteins were separated on 10% SDS-PAGE. AtlA was detected by Western immunoblotting using anti-AtlA antibodies. Representative image from at least three independent experiments is shown.

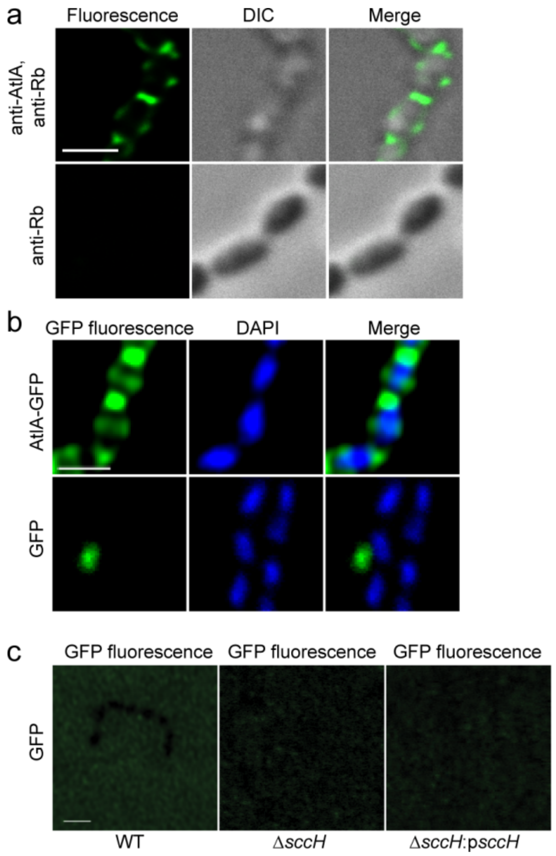

**Supplementary Fig. 7. GFP protein and fluorescent secondary antibodies do not bind intact cells of *S. mutans*, and GFP protein does not bind sacculi of *S. mutans*.**

(**a**) Cells were grown to mid-log phase, and then immunostained with anti-AtlA antibodies, followed by fluorescent secondary anti-rabbit (anti-Rb) antibodies (top panels) or fluorescent secondary anti-Rb antibodies alone (bottom panels). Cells were examined by differential interference contrast (DIC) (center panels) and fluorescence microscopy (left panels). An overlay of immunostaining and DIC is shown in the right panels. No staining was observed with fluorescent secondary anti-Rb antibodies only. The panels with the WT cells are the same as in Fig. 3 g. (**b**) Binding of AtlA-GFP (top) and GFP (bottom) to exponentially growing *S. mutans* WT. Bacterial cells show AtlA-GFP binding (top left panel), DAPI nuclear staining (center panels), an overlay of AtlA-GFP and DAPI images (top right panel), and an overlay of GFP and DAPI images (bottom right panel). No GFP binding was observed. (**c**) Binding of GFP to sacculi of WT, Δ*sccH* and Δ*sccH*:p*sccH*. No GFP signal was observed. Representative images from at least three independent experiments are shown in **a, b** and **c**. Scale bar is 1 μm.

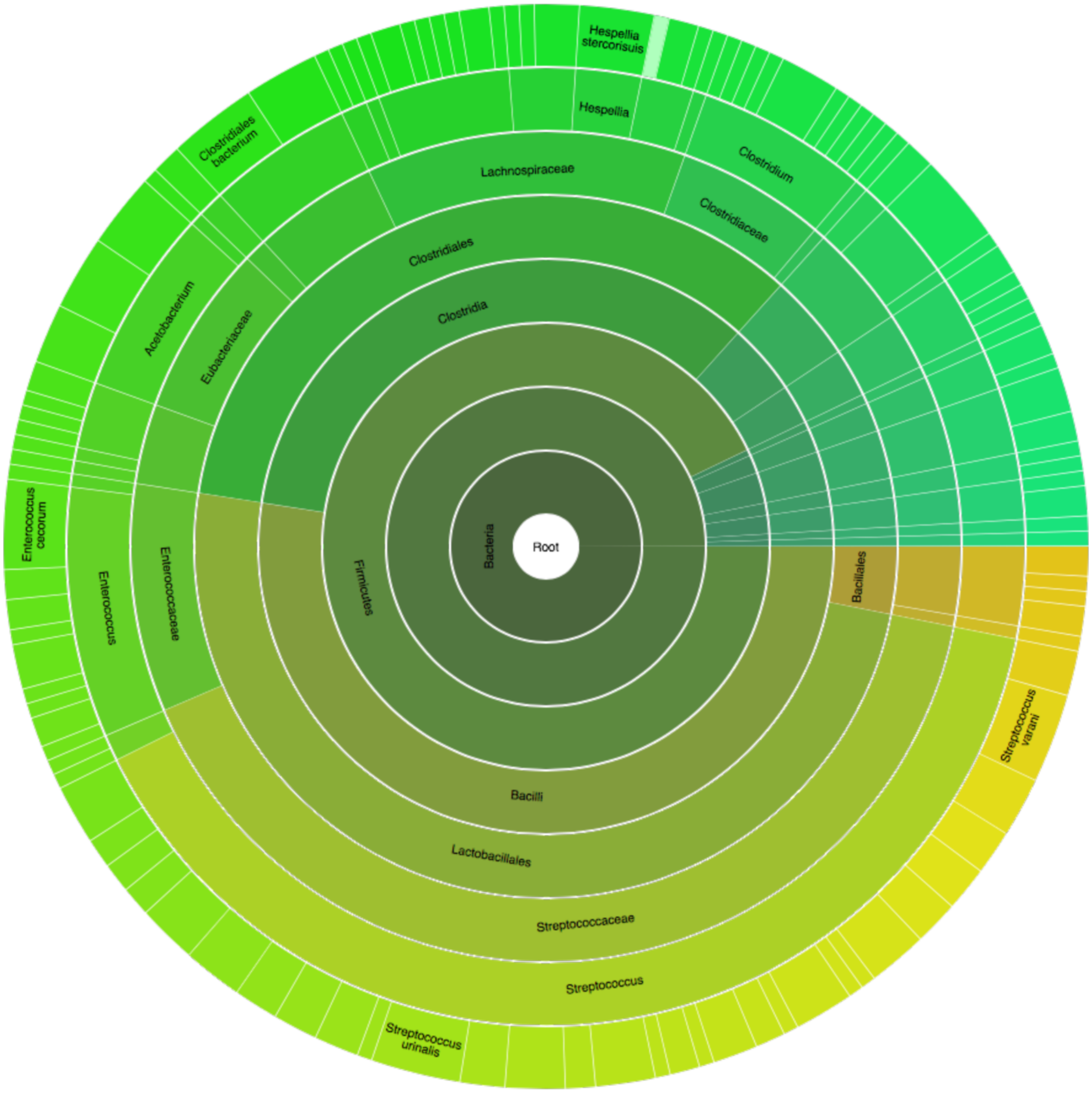

**Supplementary Fig. 8. Phylogenetic analysis of Bsp repeat domain proteins.**

A graphical representation of the distribution of Bsp domain protein family (PF08481) across bacterial species [17].

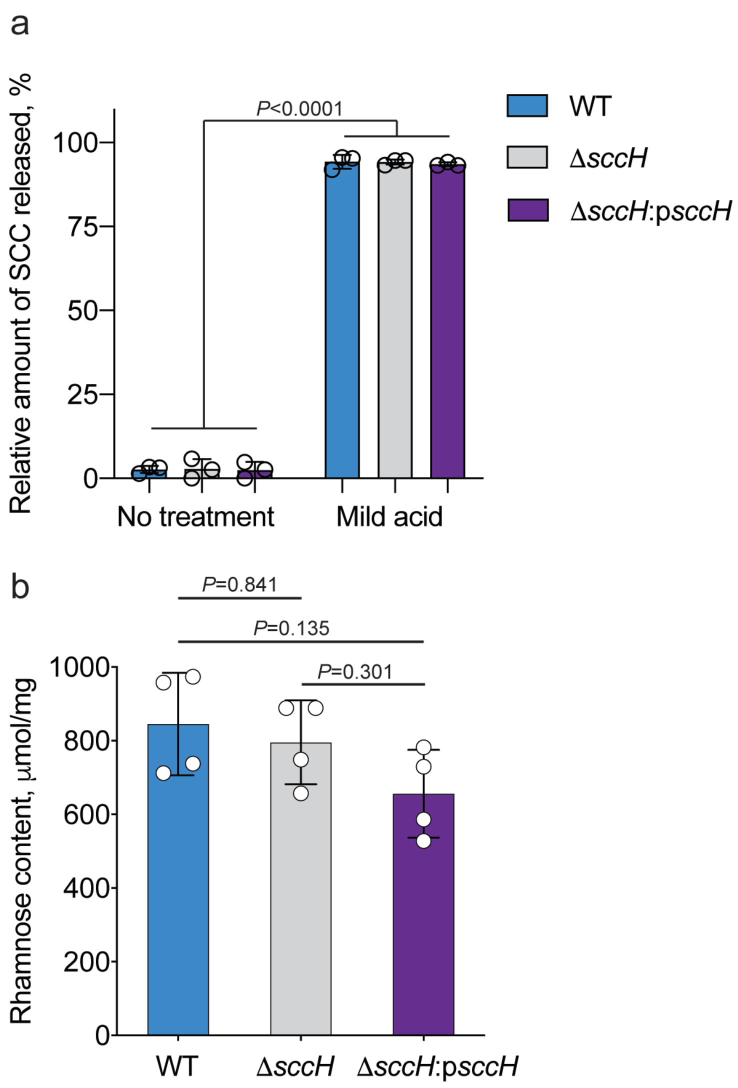

**Supplementary Fig. 9. Analysis of SCC content in cell wall of *S. mutans* WT, Δ*sccH* and Δ*sccH:psccH.***

(**a**) Release of SCCs from purified cell walls of *S. mutans* WT, Δ*sccH* and Δ*sccH:psccH* by mild acid treatment. To extract SCC from peptidoglycan, cell walls were first chemically N-acetylated as described [18] and then subjected to mild acid hydrolysis (0.02 N HCl, 100°C, 20 min) as described in Methods. The amount of SCC released from peptidoglycan was normalized to total SCC content in cell wall. (**b**) SCC content in *S. mutans* WT, Δ*sccH* and Δ*sccH:psccH*. SCCs were extracted from 1 mg of purified cell wall by mild acid hydrolysis as described in Methods. The SCC content corresponds to the Rha content which was estimated by modified anthrone assay. Columns and error bars represent the mean and S.D., respectively (biologically independent replicates in **a** and **b** are three and four, respectively). *P* values were calculated by one-way ANOVA with Tukey's multiple comparisons test. .

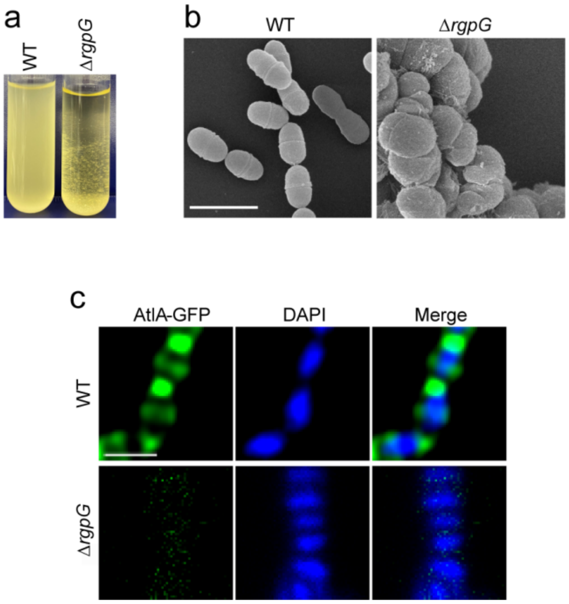

**Supplementary Fig. 9. Morphological and growth phenotypes of the Δ*rgpG* mutant.**

(**a**) The growth phenotype of WT and Δ*rgpG*. Bacteria were grown in THY broth overnight. (**b**) Scanning electron micrographs of WT and Δ*rgpG*. Bacteria were fixed, dehydrated stepwise, and viewed by scanning electron microscopy. (**c**) Binding of AtlA-GFP to exponentially growing WT and Δ*rgpG*. Bacteria were collected, fixed, incubated with AtlA-GFP and mounted on a microscope slide with mounting media containing DAPI. The WT (top) and Δ*rgpG* mutant cells (bottom) show AtlA-GFP binding (left panels), DAPI nuclear staining (center panels) and an overlay of AtlA-GFP and DAPI images (right panels, merge). The panels with the WT cells are the same as in Supplementary Fig. 7 b. The experiments depicted in **a**, **b** and **c** were performed independently three times and yielded the same results. Scale bar is 1 μm in **b** and **c**.

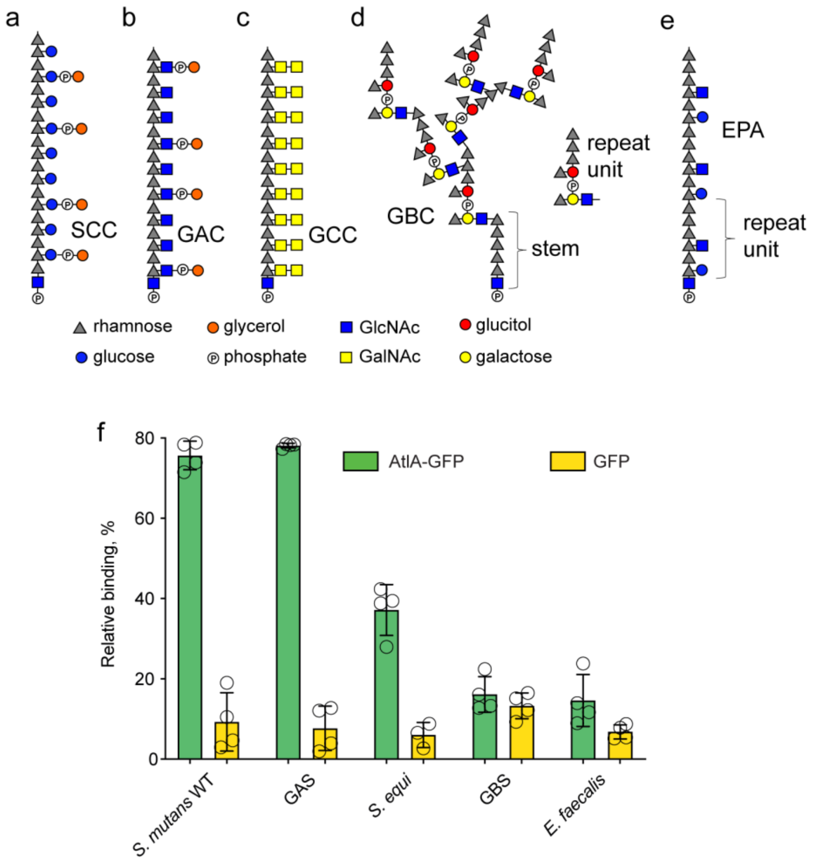

**Supplementary Fig. 10. AtlA-GFP recognizes the polyrhamnose backbone of GAC, SCC and the *S. equi* polysaccharide.**

(**a**, **b**, **c**, **d** and **e**) Schematic representation of Rha-containing cell wall polysaccharide structures. (**a**) SCC structure as determined in this study. (**b**) GAC structure as determined in [4]. Phosphate groups in SCC and GAC are involved in phosphodiester bond linking glycerol to glycosyl side-chain and polysaccharide to peptidoglycan. (**c**) *S. equi* cell wall polysaccharide (Group C Carbohydrate or GCC) as described by [19,20]. *S. mutans*, GAS and *S. equi* are the only analyzed species that contain the →3)α-Rha(1→2)α-Rha(1→ repeating backbone in their respective cell wall polysaccharide [21]. (**d**) Group B Carbohydrate (GBC) is a peptidoglycan-anchored polysaccharide in GBS. GBC is a highly branched polymer containing an →3)α-Rha(1→3)α-Rha(1→3)α-Rha(1→3)α-Rha(1→3)α-Rha(1→ unmodified pentasaccharide stem region. Additionally, →2)α-Rha(1→2)α-Rha(1→2)aRha(1→ unsubstituted Rha trisaccharides terminate the branches of the GBS polysaccharide [21-24]. Phosphate groups in GBC structure are involved in phosphodiester bond linking oligosaccharides to GBC and GBC to peptidoglycan. (**e**) Structure of the polyrhamnose backbone of *E. faecalis* cell wall polysaccharide, Epa, as determined in [25]. Epa has a repeating →2)α-Rha(1→2)α-Rha(1→3)α-Rha(1→3)α-Rha(1→2)α-Rha(1→2)α-Rha(1→ hexasaccharide backbone modified with Glc and GlcNAc side-chains. (**f**)

Co-sedimentation assay of AtlA-GFP and GFP with intact cells of *S. mutans* WT, GAS, *S. equi*, GBS and *E. faecalis*. Bacteria were grown in THY broth overnight, collected and incubated with AtlA-GFP or GFP proteins. Data is presented as a percentage of fluorescence of the pellet normalized to the total fluorescence of the sample. Columns and error bars represent the mean and S.D., respectively (n = 3 biologically independent replicates). *P* values were calculated by a two-way ANOVA with Tukey's multiple comparisons test.

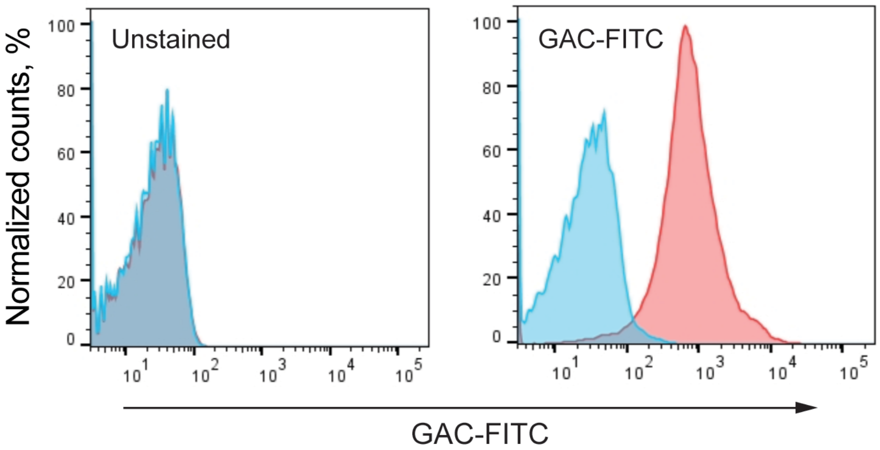

**Supplementary Fig. 11. Expression of polyrhamnose on the surface of *E. coli* PHD136.** Anti-GAC antibodies conjugated with FITC (GAC-FITC) were used to analyze polyrhamnose expression on the cell surface of *E. coli* CS2775 (parental strain) and PHD136 (CS2775 carrying pRGP1). Flow cytometry analysis confirmed that PHD136 (red) expresses polyrhamnose, whereas CS2775 (blue) does not. For flow cytometric analysis, at least 10000 events were collected. Experiments were performed independently three times and yielded the same results. Histograms of representative results are shown.

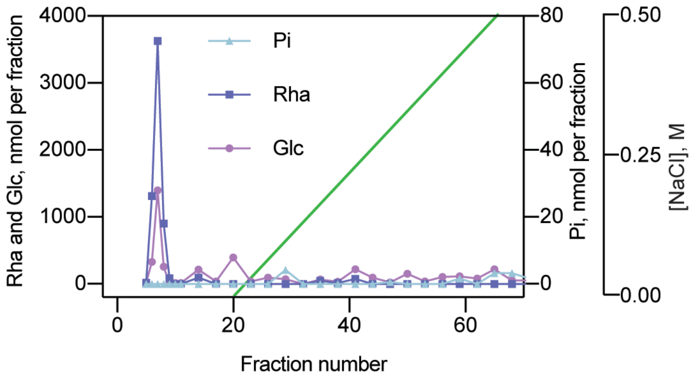

**Supplementary Fig. 12. Ion exchange chromatography of SCCs purified from Δ*sccH*.** SCC material was loaded onto Toyopearl DEAE-650M and eluted with a NaCl gradient (0-0.5 M). Fractions were analyzed for Rha and Glc contents by anthrone assay and total phosphate (Pi) content by malachite green assay following digestion with perchloric acid. The experiments were performed at least three times and yielded the same results. Data from one representative experiment are shown.

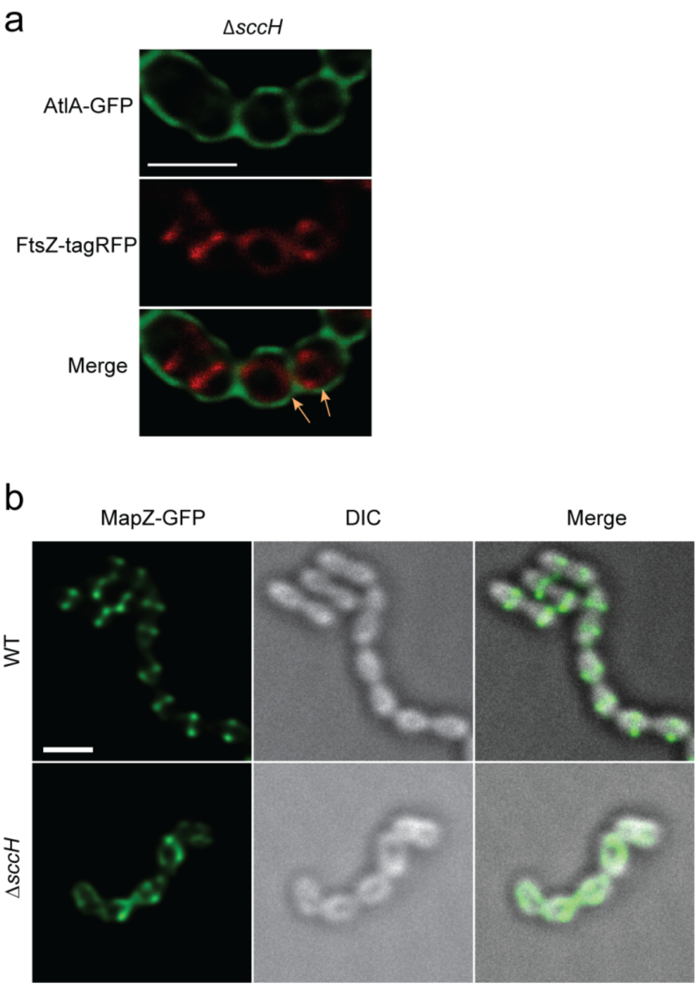

**Supplementary Fig. 13. GroP modification of SCC controls the positioning of FtsZ- and MapZ-rings.**

(**a**) Localization of FtsZ in the Δ*sccH* (strain Δ*sccH ftsZ-tagRFP*) cells. Cells expressing FtsZ-tagRFP (middle panel, red) labeled with AtlA-GFP protein (top panel, green) were analyzed. An overlay of GFP and tagRFP signals is shown in the bottom panel. Orange arrows indicate the mislocalized Z-rings. (**b**) Localization of MapZ (left panels, green) in the WT (strain *mapZ-GFP*, top panels) and Δ*sccH* (strain Δ*sccH mapZ-GFP*, bottom panels) cells. An overlay of DIC (middle panels) and GFP signal is shown in the right panels (merge). Representative images from at least three independent experiments are shown in **a** and **b**. Scale bar is 1 μm.
